# DNA methylation links lagging strand replication to transposable element control

**DOI:** 10.64898/2026.06.23.733924

**Authors:** Nora Fajri, Manon Coulée, Antoine Pigeon, Sugith Babu Badugu, Aylin Seren Güller, Susanne Bandau, Hao Jiang, Angus Lamond, Laure Ferry, Maxim Greenberg, Pierre-Antoine Defossez, Constance Alabert, Kumar Somyajit, Nataliya Petryk

## Abstract

DNA methylation is an essential epigenetic mark that silences transposable elements (TEs) in mammalian genomes^1,2^. Following DNA replication, methylation patterns must be faithfully restored^3,4^, yet how the two processes are coordinated remains unclear. Here, using strand-specific, genome-wide analyses^5–7^ in mouse embryonic stem cells, we show that DNA methylation maintenance is coupled to the lagging-strand synthesis in TE-rich regions. Paradoxically, despite this targeting, the lagging strand is more permissive for TE integration than the leading strand. Notably, insertions of full-length LINE-1s, SINEs, and satellite repeats are all enriched on the lagging strand over evolutionary time. Consequently, most TEs, particularly young elements, are oriented head-on relative to replication forks in the mouse genome, creating an unfavorable genomic configuration^8^ that is preferentially targeted by DNA methylation maintenance. Mechanistically, DNA methylation maintenance is coupled to the lagging-strand replication via UHRF1-LIG1^9^ and PCNA–PAF15^10^ interactions, and the interference in this mechanism slows Okazaki fragment maturation, and thereby potentially may facilitate TE retention. Together, we show a mechanism of TE control during DNA replication with an unexpected evolutionary interplay in which DNA methylation may facilitate, rather than solely prevent, TE expansion.

## Introduction

Eukaryotic cells must accurately replicate their chromosomes while preserving the epigenetic information embedded within chromatin components^11^. To achieve this, DNA replication forks coordinate DNA synthesis with the maintenance of chromatin features, such as histone and DNA modifications, ensuring genome and epigenome integrity across diverse genomic regions. Importantly, DNA replication is asymmetric, with the leading strand synthesized continuously and the lagging strand synthesized discontinuously as Okazaki fragments that are subsequently ligated^12,13^. This fundamental asymmetry introduces mechanistic differences in how epigenetic information is maintained, as the leading and lagging strands utilize distinct, strand-specific mechanisms for nucleosome assembly and the inheritance of histone post-translational modifications (PTMs)^14^.

In mammals, DNA methylation at CpG sites is a key epigenetic mark essential for development, genomic imprinting, and the maintenance of transcriptional states^1^. Alongside repressive histone marks, heterochromatin proteins, and silencing complexes, DNA methylation also represses the transcription of repetitive sequences derived from transposable elements (TEs), thereby maintaining genome stability and cellular homeostasis^2^.

Following each round of DNA replication, the DNA methylation patterns must be faithfully restored by remethylating the newly synthesized DNA strands at all previously methylated CpG sites to maintain symmetric methylation of both strands throughout the genome^3,4^. This maintenance is carried out by the DNA methyltransferase DNMT1 and its activator, the E3-ubiquitin ligase UHRF1 (Ubiquitin-like with PHD and ring finger domains 1)^3,4^. After replication, UHRF1 recognizes the hemimethylated DNA, where only the parental strand is methylated, and, through ubiquitination signaling, recruits and activates DNMT1 to restore full methylation^3,4^. UHRF1 ubiquitinates several substrates, including histone H3^15,16^ and the proliferating cell nuclear antigen (PCNA)-associated protein PAF15, which contains a histone-mimicking motif^17^. DNA methylation proceeds more slowly than DNA replication itself^18,19^, with distinct locus-specific kinetics across the genome, and globally faster maintenance on the lagging strands^18^. Both PAF15^17^ and H3^15,16^ ubiquitination are required for complete maintenance of DNA methylation, with PAF15 playing a predominant role during the replication-coupled phase and H3 contributing at later post-replicative stages ^17^. Efficient DNA methylation maintenance also depends on the direct recruitment of methylation factors to the replication forks^18^: UHRF1 is recruited via the histone H3–mimicking motif in the lagging strand protein DNA ligase 1 (LIG1)^9^, and DNMT1 through its PCNA-interacting motif (PIP-box)^20^. We have recently found that PAF15 is a lagging-strand-associated protein that ensures stable interaction of PCNA with chromatin during the lagging-strand synthesis^10^. Since PAF15 is a key ubiquitination substrate of UHRF1 during replication-coupled DNA methylation maintenance^17^, this observation links lagging-strand replication with the restoration of DNA methylation following DNA synthesis.

Growing evidence demonstrates that TEs exploit DNA replication forks and rely on the components of DNA replication machinery and DNA repair pathways for transposition in the genome^21–24^. Coordination of DNA replication with DNA methylation may also influence TE activity during genome duplication. However, the underlying mechanisms and biological significance of these processes remain unknown.

Here, we demonstrate that DNA methylation maintenance is coupled to lagging-strand replication at TE-rich regions. Nevertheless, the lagging strand remains more vulnerable to TE insertions and expansion than the leading strand. We further show that DNA methylation maintenance is functionally linked to lagging-strand replication via the UHRF1-LIG1-PCNA–PAF15 mechanism, suggesting that this interference may render the lagging strand susceptible to TE retention during genome duplication.

### DNA methylation maintenance is coupled with the lagging-strand synthesis at TE-regions

To investigate how DNA methylation maintenance coordinates with DNA replication genome-wide, we measured strand-specific and locus-specific binding of key DNA methylation maintenance factors UHRF1, DNMT1, and UHRF1’s interactor at the replisome, DNA ligase 1 (LIG1), to leading and lagging strands in mouse embryonic stem cells (mESCs) using SCAR-seq and OK-seq^5–7^. Analysis of average profiles around replication initiation zones (IZ) revealed that UHRF1 and DNMT1 were preferentially bound to the lagging-strand sister chromatids at leftward- and rightward-moving forks, similar to the distribution of LIG1 as well as PAF15^10^, a key activator of DNMT1^17^ and lagging-strand stability^10^ (Fig. 1a-b, Extended Data Fig. 1). Strand partitioning analysis of DNMT1, UHRF1, and LIG1 at replicated DNA revealed a significant negative correlation with replication fork directionality (RFD), confirming a genome-wide preference for the lagging strands (Fig. 1c-e). These results demonstrate that the core machinery responsible for replication-coupled DNA methylation maintenance is asymmetrically enriched at the fork with a preference for the lagging-strand replication intermediates (Fig. 1a-e, Extended Data Fig. 1). Given that a key function of DNA methylation is to silence TEs^2^, which account for ∼40 % of the mouse genome^25,26^, (Fig. 1f), we examined the genomic occupancy of UHRF1, DNMT1, and PAF15 during DNA replication and revealed that these proteins were markedly enriched at genomic regions containing TEs during DNA replication, showing a preference for Long and Short Interspersed Nuclear Elements (LINEs and SINEs) and Long Terminal Repeat retrotransposons (LTRs) (Fig. 1f-g). UHRF1 displayed a more dispersed pattern across the genome (Fig. 1f-g), consistent with its diverse roles in transcription^27^ and silencing^28^. These data suggest that DNA methylation maintenance targets the TE regions during lagging-strand replication and may reflect the epigenetic control of TEs.

**Fig. 1.**
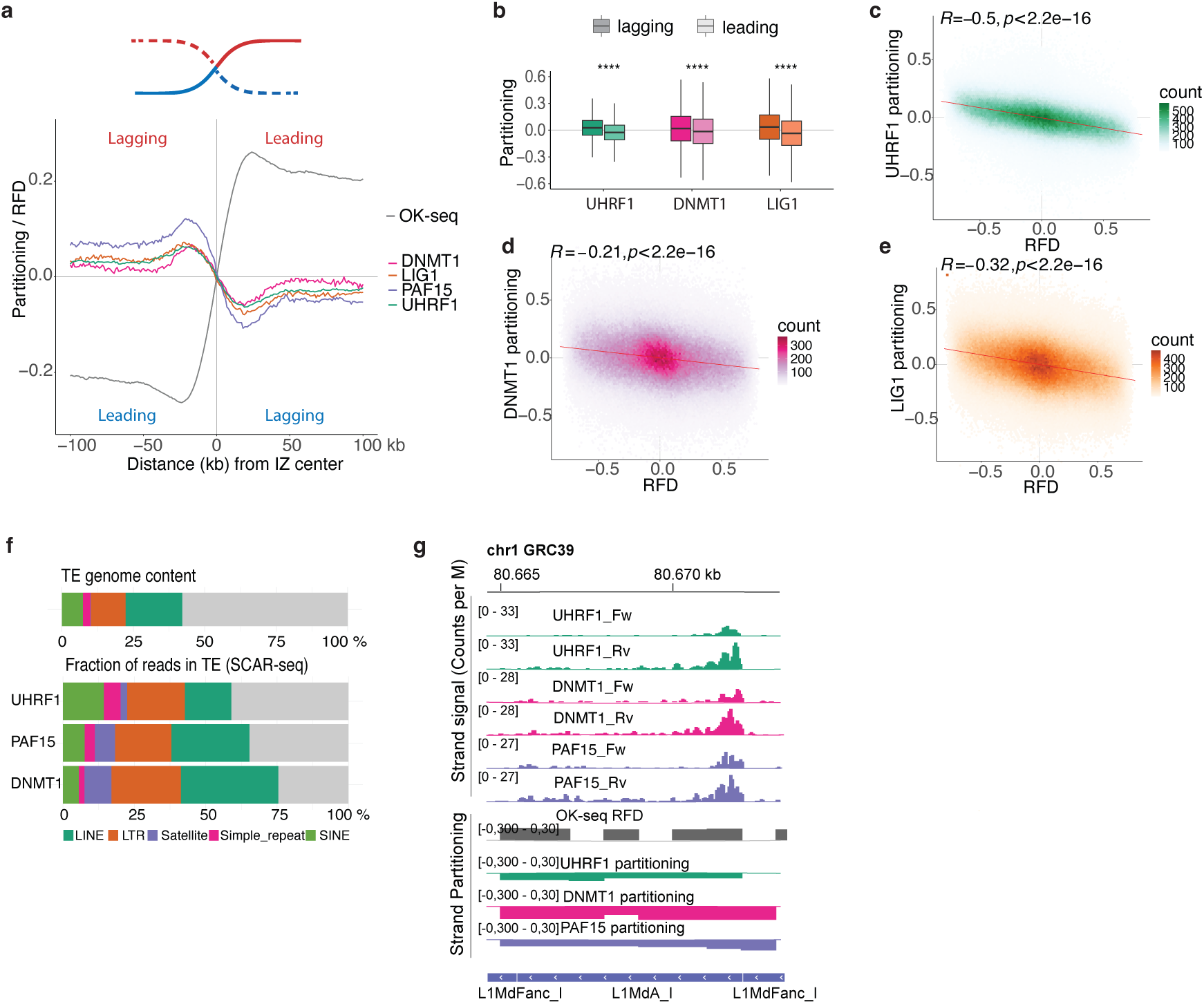
DNA methylation maintenance preferentially targets the lagging-strand in TE-containing regions. (**a**) Top: scheme indicating sister chromatids with the newly-replicated forward (red) and reverse (blue) strand switching from leading strand synthesis (full line) to lagging strand synthesis (dashed line) at initiation zones. Average profiles of RFD (grey) and UHRF1 (green), DNMT1 (pink), LIG1 (orange), and PAF15 (purple) partitioning (mean of biological replicates, shown in Extended Data Fig. 1a-c) around initiation zones, n= 4559. (**b**) Boxplot of UHRF1, DNMT1, and PAF15 partitioning values at downstream (leading, light shaded) and upstream (lagging, dark shaded) edges of initiation zones. The representation shows the mean of at least 2 biological replicates for each protein (paired Wilcoxon signed-rank test, n.s, not significant; *, p<0.05; **, p<0.01; ***, p<0.001; ****, p<0.0001). **(c,d,e)** Hexplot of UHRF1 (**C**), DNMT1 (**D**), and LIG1 (**E**) genome-wide partitioning versus RFD score. Spearman’s rank correlation coefficient is shown in the top right corner. Replicates 1, 3, and 1 are shown for UHRF1, DNMT1, and LIG1, respectively. **(f)** Top: Genomic content of TE in the mouse genome (GRCm39). Bottom: Fraction of reads in SCAR-seq of UHRF1, PAF15, and DNMT1 in TE (average of at least two biological replicates). **(g)** Genomic region representing UHRF1 (green), DNMT1 (pink), and PAF15 (purple) enrichment at forward and reverse replicated sister chromatids with their respective strand partitioning and RFD (grey).

### The lagging strand is prone to TE insertions in the genome

Given that replication-coupled DNA methylation maintenance showed a marked preference for TEs and the lagging strand during replication, we examined the relationship between TE insertions and DNA replication forks in the mouse genome. Studies in human cells have demonstrated that LINE-1 (L1) insertions are predominantly oriented head-on to the replication forks, consistent with more frequent transposition into the lagging strand during replication^21,22^. To investigate whether this phenomenon is general for other TEs, we examined the orientation of distinct TEs relative to the direction of DNA replication fork progression in the mouse genome. Notably, L1s as well as non-autonomous elements SINEs (B1, B2, B4) showed a head-on orientation bias relative to replication fork direction (Fig. 2a), indicating that these elements have transposed more frequently into the lagging strand during DNA replication in the mouse genome. Furthermore, L1s were significantly longer on average when oriented head-on to replication forks compared with those in a co-directional orientation (Fig. 2b-d). This trend is specific to L1s and was not observed for SINEs (Extended Data Fig. 2), despite SINEs utilizing L1-encoded molecular machinery and sharing a similar transposition mechanism^29,30^. The L1 mean length increased with the extent of head-on orientation, as measured by the RFD modal value (Fig. 2c), with a decline in the extreme values reflecting the global RFD distribution (Extended Data Fig. 3b). This effect was especially pronounced for evolutionarily young L1s carrying F-, A-, and N-type promoters (L1Md_Gf, L1Md_Tf, L1Md_A, L1Md_N), whereas older L1 families (Lx, L1MdV, L1Mus)^31^ showed weaker length asymmetry (Fig. 2b-c), indicating that younger elements retained full length more often when retrotransposed in a head-on orientation relative to DNA replication forks in contrast to older elements. Remarkably, the correlation between TE size and the degree of head-on orientation increased over evolutionary time (Fig. 2e-f, Extended Data Fig. 3a), suggesting that L1 improved its capacity of integration in the lagging strand. In contrast, co-directional, leading-strand-inserted L1 elements were uniformly truncated and less frequent than the lagging-strand insertions, regardless of the degree of co-orientation with the DNA replication forks and evolutionary age (Fig. 2a-b; Extended Data Fig. 3c-d).

**Fig. 2.**
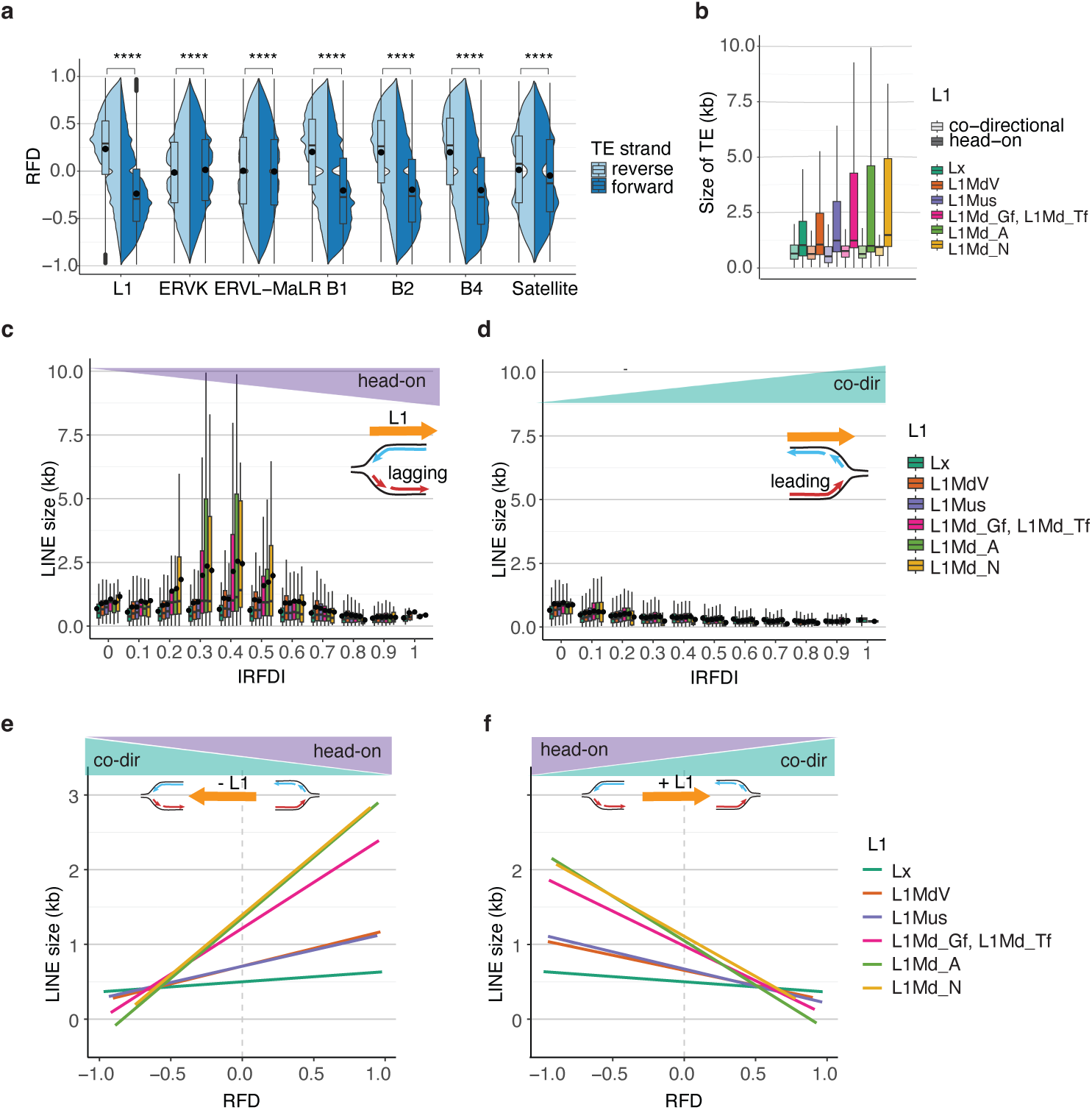
The lagging strand is permissive for TE integration. **(a)** Split violin plot showing replication fork directionality (RFD) across distinct classes of TEs, sorted by genomic orientation forward strand (dark blue) vs. reverse strand (light blue) (paired Wilcoxon signed-rank test, adjusted using Benjamini-Hochberg (BH) method, n.s, not significant; *, p<0.05; **, p<0.01; ***, p<0.001; ****, p<0.0001). **(b)** Boxplots of the length (kb) of distinct L1 families (categorised by promoter type) oriented co-directionally (light shaded) or head-on (dark shaded) to DNA replication forks. **(c, d)** Boxplots of the length (kb) of distinct L1 families (categorised by promoter type) oriented head-on **(c)** and co-directionally **(d)** to DNA replication forks, categorized by mean RFD modal values. Higher RFD mode values indicate a greater degree of head-on (or co-directional) orientation. **(e, f)** Spearman correlation between L1 size and RFD values for reverse strand (**e**) and forward strand **(f)** (individual points are represented in Extended Data Fig. 3a). Top: scheme showing the orientation (co-directional or head-on) switch relative to the DNA replication fork direction.

Strikingly, the lagging-strand-associated size insertion bias was also observed in satellite repeats, particularly for GSAT (major satellite), with centromeric regions of many chromosomes containing GSAT repeats inserted head-on relative to DNA replication forks (Extended Data Fig. 4). This indicates that lagging-strand replication is associated not only with SINEs and LINEs propagation but also with satellite expansion and may therefore contribute to the evolution of centromeres^32^.

Collectively, these findings reveal that lagging-strand replication is associated with increased insertions across major TE classes, including LINEs, SINEs, and satellite repeats in the genome over evolutionary time, and tolerates the insertions of full-length L1s. Consequently, most TEs in the mouse genome, except for LTRs, are predominantly oriented head-on toward DNA replication forks in the genome, revealing a source of potential replication-transcription conflicts^8^.

### DNA methylation maintenance targets the lagging-strand copies of head-on-oriented SINEs and L1s

Since SINEs and full-length L1s were more frequently retained in the head-on orientation to DNA replication forks in the genome, we hypothesized that head-on TEs would be preferentially targeted by replication-coupled DNA methylation maintenance. To test this, we examined how DNA methylation maintenance factors interacted with the leading- and lagging-strand copies of TEs head-on and co-directionally oriented to DNA replication forks. Indeed, all DNA methylation maintenance factors, including PAF15, UHRF1, and DNMT1, exhibited a preference for the lagging-strand copies of L1s and SINEs oriented head-on relative to DNA replication fork progression (Fig. 3a-d, S5a-c). This lagging-strand bias was not observed in co-directional L1s/SINEs or in either orientation of LTRs (Fig. 3a-c, Extended Data Fig. 5). Strikingly, DNA methylation maintenance factors were especially enriched at lagging-strand copies of evolutionarily young L1 elements (L1Md_Gf, L1Md_Tf, L1Md_A, L1Md_N), while older L1s (Lx, L1MdV, L1Mus) showed a more balanced distribution between the two strands (Fig. 3a-g). A similar pattern was observed for SINEs: DNA methylation maintenance predominantly targeted the lagging strands of young B1 SINE elements, whereas ancient SINEs, such as proto-B1, showed less strand-specific bias (Fig. S6a-c). Consistently, DNA methylation maintenance factors are also enriched at the lagging-strand copies of satellite repeats oriented head-on relative to DNA replication fork progression (Fig. S6d-f). Together, these findings demonstrate that replication-coupled DNA methylation maintenance preferentially targets lagging-strand copies of young TEs when they are oriented head-on to DNA replication forks. This strand-specific and fork-orientation-specific enrichment may therefore reflect a replication-coupled silencing.

**Fig. 3.**
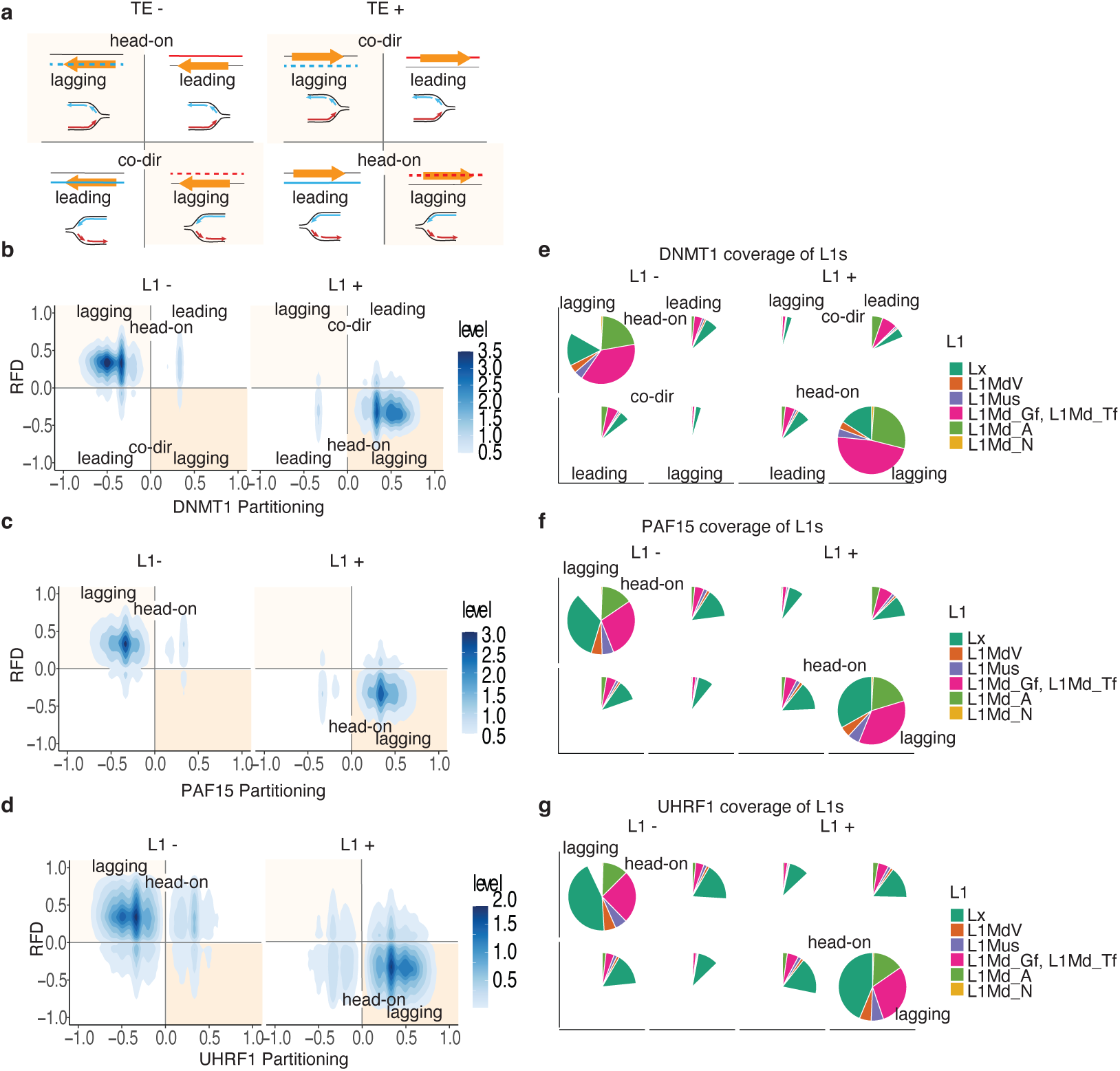
DNA methylation maintenance targets the lagging-strand copies of head-on-oriented L1s. **(a)** Scheme representing 3-dimensional separation of the replication mechanism, orientation of TE regions relative to DNA replication forks, and protein strand partitioning. The TE regions, which are predominantly replicated by the lagging strand, are highlighted in orange. Within TE in head-on orientation to DNA replication forks, the coding strand is predominantly replicated by the lagging-strand mechanism. **(b, c, d)** Density plot of partitioning of DNMT1 **(b)**, PAF15 **(c)**, and UHRF1 **(d)** within leading and lagging strand replicated copies of head-on and co-directionally oriented L1s. **(e, f, g)** Piecharts representing the coverage of distinct classes of newly-replicated copies of L1s, separated into different categories of leading and lagging replication mechanisms and relative fork orientation with DNMT1 **(e)**, PAF15 **(f)**, and UHRF1 **(g)**.

### Deficient DNA methylation maintenance leads to recruitment of H3K9me3-mediated pathways during DNA replication

Having established that DNA methylation predominantly targets TEs during lagging-strand synthesis, we focused on the interplay of DNA methylation maintenance with H3K9me3-mediated silencing pathways^2^. Previous studies have shown that the HUSH silencing complex promotes H3K9me3 deposition by SETDB1 during S phase, with a preference for the leading strands^33^. We therefore examined how H3K9me3 was distributed to the leading and lagging strands of DNA replication forks in mESCs using SCAR-seq and OK-seq^5–7^. In wild-type cells, average profiles around replication initiation zones showed the preferential enrichment of H3K9me3 at the leading-strand (Fig. 4a-b), consistent with the previous report^33^. In contrast, in *UHRF1*-KO and *LIG1*-KO cells and in the catalytically inactive mutant *DNMT1*-KO, H3K9me3 was preferentially enriched at the lagging-strand, maintaining its association with TE regions (Fig. 4a-b, S7).

**Fig. 4.**
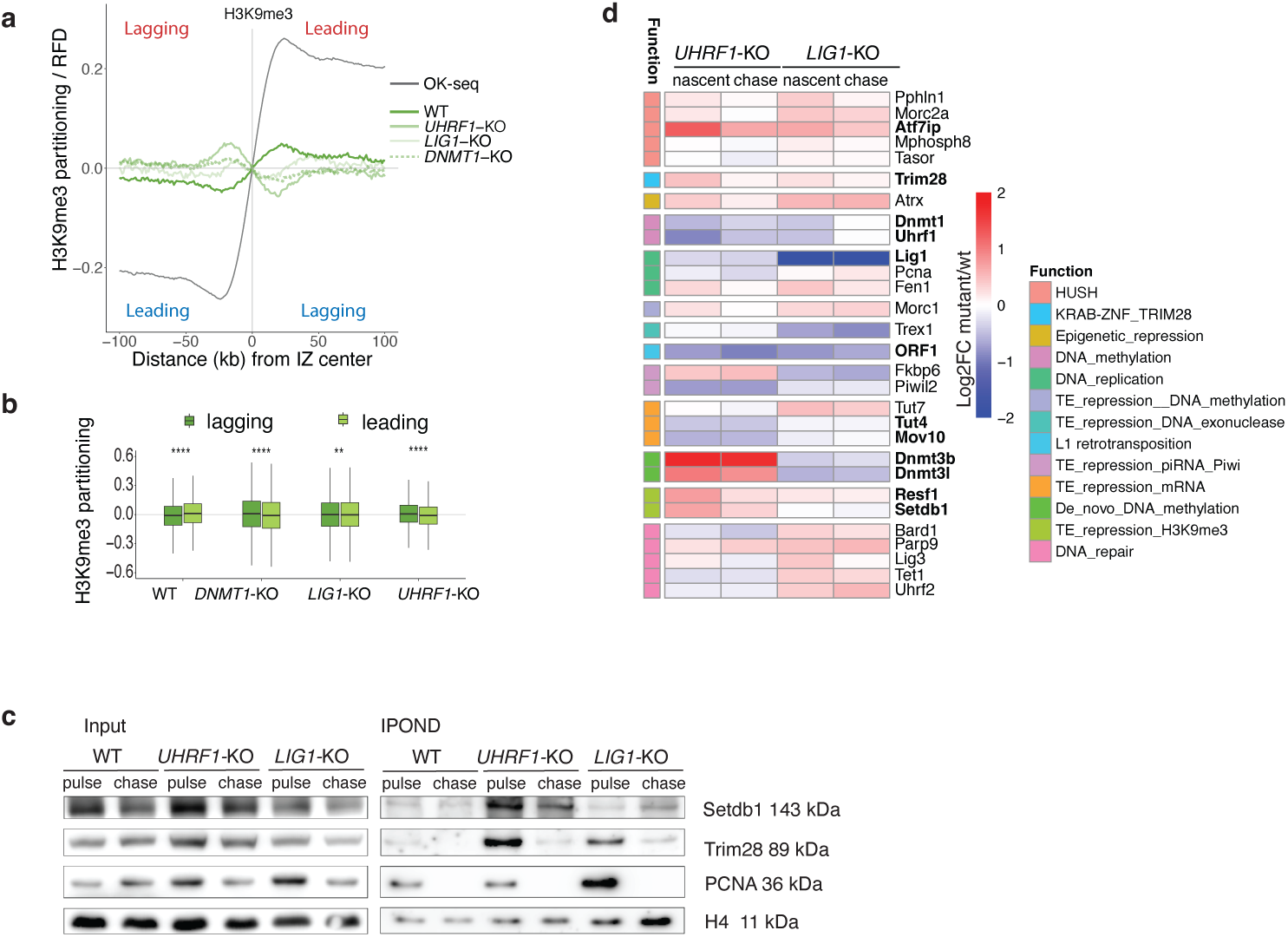
H3K9me3-mediated factors are enriched at the fork in the absence of the replication-coupled DNA methylation maintenance. **(a)** Average profiles of RFD and H3K9me3 partitioning (mean of biological replicates, represented in Extended Data Fig. 7b-e.) around initiation zones, n=4559, in wild type (WT), *UHRF1-*KO, *DNMT1-*KO, and *LIG1-*KO mutants. **(b)** Boxplot of H3K9me3 partitioning values at downstream (leading, light shaded) and upstream (lagging, dark shaded) edges of initiation zones with significant partition difference mean of replicates (paired Wilcoxon signed-rank test, n.s, not significant; *, p <0.05; **, p <0.01; ***, p<0.001; ****, p<0,0001). **(c)** IPOND western blot analysis of proteins associated with replicated DNA (iPOND) in wild type and *UHRF1*-KO or *LIG1*-KO mutants in nascent (pulse) and mature (4 h chase) chromatin. **(d)** Heatmap of log2 fold change enrichment of *UHRF1*-KO or *LIG1*-KO mutants over wild type LC-MS/MS intensities of selected candidate factors, with p-value < 0.05 at least in one condition. Factors with an adjusted *p*-value < 0.05 after Benjamini–Hochberg (BH) correction are indicated in bold. Log₂ fold change (log₂FC) values were capped at ±2.

We hypothesized that this strand shift in H3K9me3 in *UHRF1*-KO, *DNMT1*-KO, and *LIG1*-KO mutants reflects reduced UHRF1 recruitment to replication forks upon loss of replication-coupled DNA methylation maintenance. Supporting this, isolation of proteins on newly-replicated DNA (iPOND) demonstrated reduced UHRF1 recruitment to DNA replication forks in *LIG1*-KO (Extended Data Fig. 8a). Further iPOND showed the enrichment of H3K9me3 silencing factors SETDB1 and TRIM28 at DNA replication forks (pulse) compared to post-replication (chase) in *UHRF1*-KO and *LIG1*-KO (Fig. 4c). Moreover, iPOND-mass spectrometry analysis confirmed increased presence of key factors involved in H3K9me3-mediated silencing of TEs^34^, including components of the HUSH complex (Mpp8 and Periphilin), MORC2, and SETDB1’s chaperone ATF7IP in both *UHRF1*-KO and *LIG1*-KO (Fig. 4d, Extended Data Fig. 8b-e). These data demonstrate that, in the absence of replication-coupled DNA methylation maintenance, H3K9me3-mediated mechanisms are actively recruited, suggesting a potential interplay between these silencing pathways to preserve TE repression during DNA replication.

### UHRF1 functionally couples DNA methylation maintenance to lagging-strand replication through the PCNA–PAF15 axis

To test whether perturbation of DNA methylation maintenance can affect lagging-strand replication, we investigated the functional connection between these two processes through the UHRF1-LIG1^9^ and PCNA–PAF15 axis^10,17^. We treated human osteosarcoma U2OS cells with the DNMT1 inhibitor 5-aza-2′-deoxycytidine (aza-dC). Using quantitative image-based cytometry (QIBC) and QIBC-proximity ligation assays (QIBC-PLA), we found that aza-dC treatment increased PCNA retention on chromatin and stabilized PCNA–DNMT1 interactions in a PAF15-dependent manner (Fig. 5a, Extended Data Fig. 9a-b), indicating that stalling of DNA methylation maintenance affects PCNA dynamics at the replication fork through PAF15. Consistently, DNMT1 mutant and triple DNMT1/3A/3B mutants abolished UHRF1-dependent PAF15 ubiquitination and reduced PAF15 protein levels in mESCs (Fig. 5b), supporting that DNA methylation is required to maintain the PCNA-PAF15 regulatory axis. Furthermore, UHRF1 knockdown markedly reduced PCNA interactions with both DNMT1 and the core PCNA-interacting lagging-strand synthesis factors FEN1 and LIG1^12^, without affecting PCNA loading on chromatin (Fig. 5c, Extended Data Fig. 9c-h), demonstrating that UHRF1 regulates both DNA methylation maintenance and lagging-strand replication through PCNA.

**Fig. 5.**
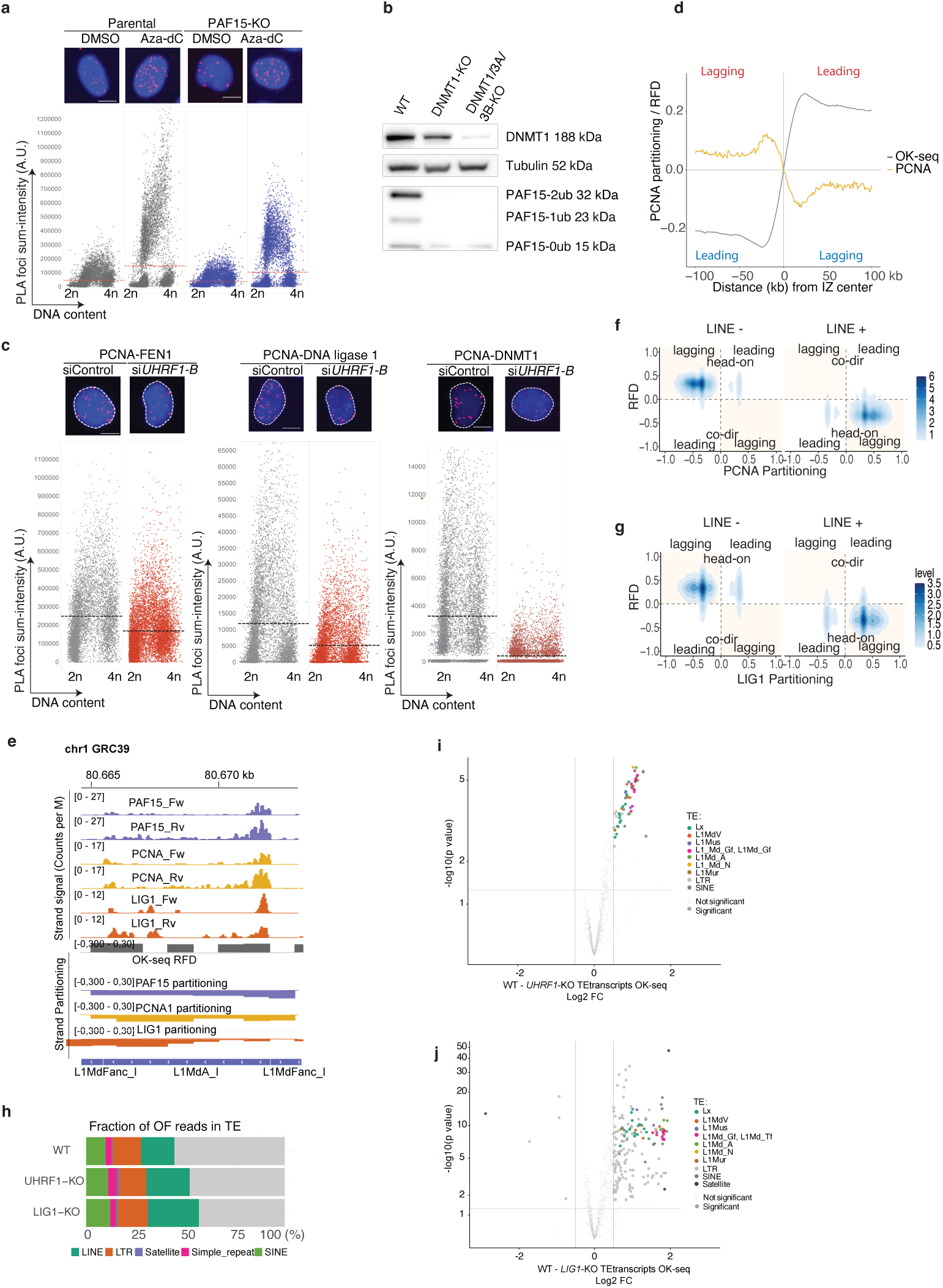
Replication-coupled DNA methylation maintenance at transposable elements interferes with the PCNA turnover during the lagging-strand synthesis. **(a)** Representative PLA images and QIBC analysis of PCNA–DNMT1 PLA foci in parental and PAF15 knockout cells treated with decitabine (∼8,000 cells analyzed per condition). DNA content: 2N (G1 phase), 4N (G2 phase). **(b)** Whole cell western blot of PAF15 ubiquitination state in *DNMT1*-KO, *DNMT1/3A/3B*-KO cells. **(c)** Representative PLA images and PLA-QIBC analysis of PCNA interaction with FEN1, LIG1, and DNMT1 in wild-type and UHRF1-depleted U2OS cells. **(d)** Average profile of RFD and PCNA partitioning (mean of 3 replicates) around initiation zones, n= 4559. **(e)** Genomic example of PCNA (yellow) and LIG1 (orange) strand-specific enrichment during DNA replication and partitioning. **(f, g)** Density plot of partitioning of PCNA **(f)** and LIG1 **(g)** within leading and lagging strand replicated copies of head-on and co-oriented LINEs. **(h)** Fraction of Okazaki fragments reads within TE in wild type, *UHRF1*-KO, and *LIG1*-KO cells (average of 2 replicates). **(i, j)** Volcano plots showing differential enrichment of OK-seq reads within transposable element (TE) families in UHRF1-KO and LIG1-KO mutants relative to wild-type cells. The x-axis indicates the log2 fold change in TE-associated OK-seq read counts, and the y-axis shows the −log10 *P* value. TE families exhibiting significant differential enrichment are shown as filled circles. TE families with an adjusted *P* value (p.adjust) < 0.05 were considered significant.

To determine whether this functional connection manifests at TEs specifically, we assessed strand- and locus-specific PCNA occupancy during replication using SCAR-seq and OK-seq^5–7^. Consistent with previous findings in yeast^35^, PCNA occupancy was higher at the lagging strand than the leading strand genome-wide (Fig. 5d, Extended Data Fig. 10a-b). Strikingly, PCNA occupancy was not uniform: PCNA was markedly enriched at TEs (Fig. 5e, Extended Data Fig. 10e), revealing a strong preference for lagging-strand copies of head-on-oriented L1s (Fig. 5g) and mirroring the distribution of DNMT1, PAF15, and UHRF1 (Fig. 3). Similarly, LIG1 was enriched at lagging-strand copies of L1s (Fig. 5e-f) and, to a lesser extent, SINEs (Extended Data Fig. 10c-d). Together, this co-enrichment of PCNA and LIG1 at lagging-strands in TE regions suggests slower Okazaki fragment maturation when replication forks traverse TE regions.

To further reveal if UHRF1 and LIG1 could regulate Okazaki fragment maturation in TE regions, we compared DNA replication dynamics and Okazaki fragment maturation using OK-seq^7,36^ in *LIG1*-KO and *UHRF1*-KO cells. RFD profiles were broadly preserved, and initiation zone locations largely overlapped between wild-type cells and *LIG1*-KO and *UHRF1*-KO conditions (Extended Data Fig. 11a-b), indicating no major disruption of the overall replication program. Nevertheless, both *LIG1*-KO and *UHRF1*-KO knockouts increased the fraction of Okazaki fragment reads mapping within TEs, especially within young families of L1s (L1Md_A, L1Md_Gf/L1Md_Tf and L1Md_N) (Fig. 5f-j), the elements that display preferential head-on orientation to replication forks and enrichment of UHRF1 and other DNA methylation maintenance factors at their lagging-strand copies. This is consistent with slower lagging-strand maturation in TE-containing regions when either DNA methylation maintenance or lagging-strand replication is perturbed.

Collectively, these data demonstrate that replication-coupled DNA methylation maintenance and lagging-strand synthesis are functionally linked through the UHRF1-LIG1-PCNA–PAF15 mechanism, and that perturbation of either process impairs lagging-strand replication within TE-rich regions. We propose that this interference may create a window for TE integration machinery to engage the lagging strand, rendering it particularly susceptible to TE insertions, and may underlie the preferential retention of TEs in the mammalian genome.

## Discussion

We describe a strand-specific mechanism of TE control that intrinsically coordinates the lagging-strand DNA replication with DNA methylation maintenance. Key proteins involved in DNA methylation maintenance were enriched on the lagging strands of DNA replication forks in TE-containing genomic regions, revealing replication-coupled targeting of TEs by DNA methylation maintenance. However, the lagging strand appeared to be more prone to TE insertions, resulting in a global head-on orientation of major TE classes relative to DNA replication forks, including SINEs, satellites, and intact L1s. Because SINEs depend on L1-encoded proteins, the insertion bias observed for both L1s and SINEs likely reflects a common replication-dependent mobilization pathway for autonomous and non-autonomous retrotransposons through preferential targeting of the lagging strand. This appeared to be an effective evolutionary strategy, and half of the mammalian genome is composed of SINEs and L1s.^37^ The lagging-strand inserted SINEs and L1s may be better tolerated by the host since they can be effectively silenced by replication-coupled DNA methylation maintenance, while TEs exploit the discontinuous synthesis for their propagation. A possible mechanistic explanation may arise from the interaction between L1-encoded reverse transcriptase ORF2 and PCNA^38^. The lagging strand is characterized by discontinuous synthesis and iterative PCNA loading during Okazaki fragment maturation^12,13^, which may provide windows for ORF2 engagement^38^ as well as preferred substrates for target-primed reverse transcription^39^, thereby promoting TE insertions in head-on orientation relative to replication fork progression. In contrast, PCNA on the leading strand is more stably integrated within the replisome and with DNA polymerase epsilon, making it inherently less accessible^40,41^.

Remarkably, we show that the bias towards preferential lagging-strand insertions of L1 elements increases over evolutionary time. Given that DNA replication timing and, therefore, fork directionality are highly conserved features across evolution, reflecting large-scale genome organization^42^, this suggests that the L1 retrotransposition machinery may have progressively increased its affinity for the lagging strand, with PCNA–PAF15 playing an important role in this interplay. Consistent with this, we demonstrate that the lagging-strand copies of younger elements are strongly marked by DNA methylation maintenance and PCNA-dependent lagging-strand replication factors. Furthermore, the lagging-strand replication is functionally coupled with DNA methylation maintenance through PCNA-PAF15, and these processes mutually influence one another.

The strand specificity of TE insertion also has implications for how host defense mechanisms operate beyond epigenetic silencing. DNA repair plays a critical role in host defense against TEs, including 5’ truncation of L1s during transposition as a protective mechanism^22,43^. Key repair proteins belonging to homologous recombination pathways inhibit L1 retrotransposition^23,24^. Our findings suggest that L1 truncation may be more frequent during leading-strand synthesis than during lagging-strand synthesis. This may be related to differences in DNA repair efficiency between the strands, although this remains to be directly tested, potentially contributing to the emergence of strand-specific epigenetic strategies in host defense against TE activity. Therefore, as a cost of discontinuous synthesis, the lagging strand is associated with a higher frequency of TE insertion events and increased retention of full-length L1, compared to the leading strand, with potential implications for genome stability and evolution.

The host genome deploys replication-coupled DNA methylation as a control strategy for TE silencing at the lagging strand. DNMT1-UHRF1 recruitment to replication forks and its regulation via PAF15-PCNA-LIG1 position the DNA methylation maintenance machinery preferentially at nascent lagging-strand synthesis, potentially ensuring restoration of DNA methylation–mediated silencing after DNA synthesis^18^. At the same time, competition between UHRF1 and SETDB1 for H3K9me3 binding^44^ may transiently delay histone-based^33^ repression during fork passage at the lagging strand. The lagging strand may therefore operate under a distinct timing of epigenetic regulation compared to the leading strand, in which DNA methylation precedes H3K9me3 during S phase, through UHRF1 activity. This strand-specific control may be important for stability of the PCNA-PAF15 regulatory axis and accurate lagging-strand replication through TE-rich regions.

Over evolutionary time, the lagging-strand retrotransposition bias may have broader consequences for genome architecture and stability. Tolerance of the lagging-strand for TE insertions may be associated with genome expansion^45^. Beyond evolution, the preferential retention of full-length transcriptionally competent L1s in head-on orientation relative to DNA replication forks increases the potential for replication-transcription conflicts^8^, with direct implications for genome stability. Additionally, L1 activity has been broadly linked to inflammatory disorders and cancer^43,46^, and failure of the replication-coupled silencing may contribute to pathological TE reactivation. Finally, the coordination between DNA replication and DNA methylation maintenance may also regulate L1 activity during epigenetic reprogramming in early development, a context in which both processes are dynamically remodeled^47–49^.

Together, our results reveal a strand-asymmetric mechanism of TE control during DNA replication, in which the lagging strand is targeted for replication-coupled DNA methylation maintenance and at the same time, reveals preferential insertion of TEs. The PCNA–PAF15–UHRF1 axis coordinates both Okazaki fragment maturation and DNA methylation maintenance, but also renders them mutually vulnerable: perturbation of either impairs the other, potentially creating conditions that favor TE retention. This mechanistic coupling may partly explain why mammalian genomes have accumulated such a high TE burden despite robust epigenetic defenses (Extended data Fig. 12).

## Acknowledgments

We acknowledge the sequencing expertise of the I2BC High-throughput sequencing facility, supported by France Génomique and funded by the French National Program “Investissement d’Avenir” ANR-10-INBS-09.

## Author contributions

N.P. designed and supervised the study. N.F., A.P., and N.P. performed most of the experiments. M.C. performed genomic analyses and visualizations under the supervision of N.P. H.J. performed mass-spectrometry analyses under the supervision of A.L.; S.B.B and A.S.G. performed QIBC and QIBC PLA experiments under the supervision of K.S. S.B. performed proteomic data normalization and DEG analysis under the supervision of C.A.; L.F. performed the iPOND western blot experiment in *LIG1*-KO cells under the supervision of P-A. D; M.G. generated and validated the *DNMT1*-KO cell line. N.P. prepared figures and wrote the manuscript with input from all co-authors. All authors read and approved the final version.

## Extended data

**Materials and Methods**

**Extended data Figs. 1 to 12.**

**Table 1.** List of the 7139 proteins differentially associated with replicated chromatin.

**Table 2.** Genomic positions of replication initiation zones in mESC wild type and *UHRF1*-KO and *LIG1*-KO mutants.

