## Extended data and methods for "DNA methylation links lagging strand replication to transposable element control"

**This PDF file includes:**

**Materials and Methods.**

**Extended data Figs. 1 to 12.**

**Table 1.** List of the 7139 proteins differentially associated with replicated chromatin.

**Table 2.** Genomic positions of replication initiation zones in mESC wild type and UHRF1-KO and LIG1-KO mutants.

**References (50-73).**

### Materials and Methods

#### Cell culture

E14Tg2a mouse embryonic stem cells (mESCs) were cultured in standard ESC medium, consisting of Dulbecco's Modified Eagle Medium (DMEM, Gibco) supplemented with 15 % KnockOut™ Serum Replacement (KSR, Gibco), 1 mM sodium pyruvate (Gibco), 1× non-essential amino acids (Gibco), 1× Penicillin-Streptomycin-L-Glutamine (Life Technologies), 10 ng mL$^{-1}$ mouse leukemia inhibitory factor (LIF, Miltenyi), and 0.1 mM β-mercaptoethanol (Sigma). E14 LIG1-KO and UHRF1-KO mutants were obtained from[9,50]. U2OS cells were cultured and siRNA transfection was performed as previously described[10].

#### Genome editing

*DNMT1*-KO cells were generated in E14Tg2a using gRNAs that were previously described[51]. The gRNA targeting the DNMT1 gene was cloned into pX459 (Addgene plasmid #48139) and transfected into E14Tg2a cells. After puromycin selection, ~96 colonies were picked and screened by DNA digestion with the DNA methylation-sensitive restriction enzyme HpaII. DNA methylation mutant clones were further confirmed by Sanger sequencing. For the *DNMT1*-KO clone used in this study, loss of DNA methylation was confirmed by the luminometric methylation assay (LUMA) [52], which was performed exactly as described[53].

#### Western blot

1 × 10$^{6}$ cells were cultured in 6-well plates overnight, washed with 1× PBS, and lysed on ice for 5 min in 50 µL RIPA buffer (25 mM Tris-HCl, 150 mM NaCl, 1 % NP-40, 1 % sodium deoxycholate, 0.1 % SDS) supplemented with cOmplete™ protease inhibitor (Roche). Lysates were sonicated (30 s on/30 s off cycles for 1 min 30 s at 60 % amplitude), mixed with Laemmli loading buffer containing 50 mM DTT, and denatured at 95 °C for 10 min.

Proteins were separated on 4–12 % Bis-Tris gels using 1× MOPS buffer (120 V, 1 h) and transferred to nitrocellulose membranes using Tris-glycine transfer buffer containing 10 % ethanol (120 V, 400 mA, 1 h 20 min, on ice). Membranes were blocked in 5 % milk in PBST (0.05 % Tween-20) for 1 h at room temperature and incubated overnight at 4 °C with primary antibodies: H4 (16047-1-AP; 1:600), PAF15 (sc-390515; 1: 500), PCNA (sc-56; 1: 500), Tubulin (66031-1-Ig; 1: 1000), UHRF1 (sc-373750; 1: 500), LIG1 (MA1-23189; 1: 1000), DNMT1 (ab188453; 1: 1000), TRIM28 (KAP1) (ab10483; 1: 1000), and SETDB1 (ab313862; 1: 1000).

Membranes were washed three times with PBS-T (0.05 % Tween-20), incubated with HRP-conjugated secondary antibodies (anti-mouse ab6789, 1: 5000; anti-rabbit ab6721, 1: 10000) for 1 h at room temperature, and developed using SuperSignal™ Pico Plus substrate. Images were acquired using an Amersham Imager 680.

#### SCAR-seq (Sister Chromatid After Replication sequencing)

SCAR-seq was performed as previously described [5,10,54] with the following modifications.

***CUT&RUN SCAR-seq:*** For UHRF1 SCAR-seq, chromatin immunoprecipitation (ChIP) was replaced by the CUT&RUN method which was performed as previously described[55]. Cells were grown to 80 % confluence in 15 cm dishes and labeled with 20 µM EdU for 20 min. Cells were then washed with wash buffer (20 mM HEPES, 150 mM NaCl, 0.5 mM spermidine (S0266, Sigma), and 1× cOmplete protease inhibitor (11873580001, Roche), bound to concanavalin A–coated beads (86057-3, Clinisciences), and permeabilized with digitonin (CHR103, Merck Millipore; 0.05 % final concentration in wash buffer).
Samples were incubated with 2 µg of anti-UHRF1 antibody (Santa Cruz, sc-373750) for 1 h at room temperature, followed by three washes with the digitonin wash buffer (dig-Wash buffer, 0.05 %). Samples were then incubated with protein A–micrococcal nuclease (2.5 ng of pA-MNase in dig-wash buffer) for 1 h at room temperature and washed again with dig-Wash buffer.
MNase was activated by the addition of 2 mM $CaCl_2$, resulting in cleavage of DNA proximal to antibody-bound proteins. DNA fragments were released into the supernatant following incubation at 37 °C for 20 min with shaking and subsequently collected for downstream steps including click-biotinylation, library preparation, and purification of newly synthesized strands, as previously described[10,54].
***X-SCAR-seq:*** For DNMT1, LIG1, and PCNA, SCAR-seq was performed with the chromatin immunoprecipitation step (ChIP), as previously described[10]. Briefly, $60 \times 10^6$ cells were labeled with EdU for 20 min and crosslinked with 1 % formaldehyde for 10 min at room temperature. Crosslinking was quenched by the addition of 0.1 M glycine for 5 min. Cells were washed twice with ice-cold PBS, collected by scraping, and pelleted by centrifugation.
Cell pellets were lysed for 20 min at room temperature in lysis buffer consisting of a mixture of 1/3 dilution buffer (100 mM Tris-HCl pH 8.6, 100 mM NaCl, 5 mM EDTA, 5 % Triton X-100) and 2/3 SDS buffer (50 mM Tris-HCl pH 8.1, 100 mM NaCl, 5 mM EDTA, 0.5 % SDS), supplemented with cOmplete protease inhibitor.
Chromatin was sonicated using a Diagenode Bioruptor Pico to obtain DNA fragments of approximately 250–300 bp (30 cycles: 30 s ON / 30 s OFF) in 15 ml tubes.
For each ChIP, ~300 µg of chromatin (based on DNA concentration) was incubated with 10 µg of antibody against DNMT1 (Abcam, ab188453), LIG1 (Invitrogen, MA1-23189), or PCNA (Santa Cruz, sc-56). Immune complexes were captured using anti-mouse or anti-rabbit IgG Dynabeads (Thermo Fisher, 11201D and 11203D, respectively). DNA–protein complexes were reverse crosslinked at 55 °C for 4 h. An aliquot corresponding to 1 % of the lysate was reserved as input.
For both methods, click-mediated biotinylation of newly replicated DNA, nascent strand isolation, and library preparation were performed as previously described [5,10,54] using xGen UDI-UMI Adapters (IDT). Libraries were sequenced in paired-end mode on an Illumina NextSeq 2000.
Histone SCAR-seq was performed as previously described [5,54] using an antibody against H3K9me3 (Abcam, ab176916).

**SCAR-seq data processing**

Raw reads were trimmed and filtered for quality (Q < 20) using Cutadapt (v. 2.6), then aligned to the mouse reference genome (GRCm39.109) with bowtie2 (v. 2.4)[56]. . Duplicate reads were marked and removed using Picard MarkDuplicates function (v. 3.4) (RRID:SCR_006525).. Processed BAM files were separated into forward and reverse strands based on SAM flags using SAMtools (v. 1.13)(forward: -f 83 and -f 163; reverse: -f 99 and -f 147), with flags adjusted to account for the inverted orientation of IDT UDI-UMI adapters.

Read coverage was calculated in 1 kb bins using multiBamSummary (deepTools, v. 3.5.4) [57], and bins with CPM ≤ 0.3 were excluded from downstream analyses. Strand-specific partitioning for PAF15, PCNA, DNMT1, LIG1, UHRF1, and H3K9me3 was computed from binarized mapped reads as:

$$\text{Partitioning} = \frac{F - R}{F + R}$$

where F and R correspond to the number of reads mapped to the forward and reverse strands, respectively.

**OK-seq**

OK-seq was performed as previously described[7,36] with modified adapter sequences:

**Adapter1w-mixA:** [SpC3] ACA CTC TTT CCC TAC ACG ACG CTC TTC CGA TCT

**Adapter1c-mixA:** [SpC3] NNNNNN AGA TCG GAA GAG CGT CGT GTA GGG AAA GAG TGT

**Adapter2w-mixA:** [Phos] AGA TCG GAA GAG CAC ACG TCT GAA CTC CAG TCA [SpC3]

**Adapter2c-mixA:** [SpC3] TGA CTG GAG TTC AGA CGT GTG CTC TTC CGA TCT NNNNNN [SpC3]

Libraries were sequenced in paired-end mode on an Illumina NextSeq 2000.

**OK-seq data processing**

Raw reads were trimmed and filtered for quality (Q < 20) using Cutadapt (v. 2.6), then aligned to the mouse genome (GRCm39.109) with bowtie2 (v. 2.4) [56]. Duplicate reads were marked and removed using Picard MarkDuplicates function (v. 3.4) (RRID:SCR_006525). Processed BAM files were split into forward and reverse strands based on SAM flags using SAMtools v1.13(*21*) (forward: -f 99 and -f 147; reverse: -f 83 and -f 163).

Read coverage was computed in 1 kb bins using multiBamSummary (deepTools, v. 3.5.4) [57], and bins with CPM ≤ 0.3 were excluded. Replication fork directionality (RFD) was calculated as:

$$\text{RFD} = \frac{R - F}{R + F}$$

where R and F represent the number of reads mapped to reverse and forward strands, respectively. Initiation zone (IZ) positions were identified as previously described[10], considering only regions with $RFD_{max} > 0$ and $RFD_{min} < 0$. IZs were retained according to their size (≥10 kb), efficiency >10% (ΔRFD ≥ 0.2), and must have  minimal 50 % overlap between biological replicates (BEDtools intersect, v. 2.30.0 [58], yielding 4559 regions in wild-type cells, 9120 in UHRF1-KO mutant, and 3098 in LIG1-KO. Spearman correlation was performed on the RFD score between mutant and wild-type conditions using the plotCorrelation function (deepTools, v. 3.5.4), and the common IZ position was shown using the UpSetR package.

**SCAR-seq and OK-seq data analysis**

SCAR-seq partitioning and RFD were smoothed using a 30 kb sliding window (±15kb), and extreme values are filtered out (|RFD| > 0.00001 and |RFD| < 0.99999). Average profiles were computed according to a 1-kb bin partition and RFD distance from IZ (IZ ±100kb). Differences in SCAR-seq partitioning between the left and right ends of IZs were assessed using a paired Wilcoxon signed-rank test on at least two independent biological replicates.

Genome-wide Spearman's rank correlations were calculated in 1 kb bins and visualized as hexplots. Heatmaps of SCAR-seq profiles around IZs (±0.2 Mb) were generated using deeptools (v. 3.5.4) plotHeatmap function[57]. Genomic regions were visualized with IGV viewer.

**Transposable elements analysis**

Transposable elements (TEs) were generated using RepeatMasker[59] from UCSC Repeat Browser on the mouse genome (mm39/GRCm39). SCAR-seq and OK-seq strand-specific reads were quantified in TEs using deepTools (v. 3.5.4) [57] multiBamSummary, and SCAR-seq partitioning and OK-seq RFD were calculated as for genome-wide analyses.

SCAR-seq partitioning at TE regions distinguishes sister chromatids based on the orientation of newly replicated DNA strands relative to TE orientation: the "old strand" corresponds to the sister chromatid carrying TE coding sequence at the parental strand (opposite to SCAR-seq read direction), while the "new strand" corresponds to the sister chromatid carrying TE coding sequence at the newly synthesized strand (same direction as SCAR-seq reads). OK-seq RFD values indicate replication fork direction relative to TE transcription: "head-on" corresponds to forks moving opposite to the TE orientation (RFD/TE -/+ or +/-), whereas "co-directional" corresponds to forks moving in the same direction as the TE orientation (RFD/TE -/- or +/+).

Statistical analyses (violin plots and boxplots) were performed using a Wilcoxon test, adjusted by the Benjamin-Hochberg test, across at least two independent biological replicates. LINE L1 and SINE B1 elements were further classified into six subcategories according to their respective evolutionary ancestries[31,60].

Density plots were generated according to the mapped TE genomic orientation and its respective RFD value and partition score calculated within TE.

**Okazaki fragment quantification within TEs**

Enrichment of Okazaki fragments within transposable element (TE) regions was quantified using TEtranscripts[61]on OK-seq mapped BAM files before the removal of multimapping reads, using default TEtranscripts parameters. Okazaki fragment reads were assigned to TE families based on the RepeatMasker annotation. Low enriched TE are filtered out (CPM > 0.2 in at least 2 samples).

Differential TE enrichment between wild-type (WT) and mutant samples was assessed using DESeq2 (v1.42.1). Two independent biological replicates were analyzed for WT samples, and at least three independent biological replicates for each mutant genotype. TEs with an adjusted *P* value (p.adjust) < 0.05 were considered significantly differentially enriched.

**IPOND**

IPOND experiments were performed essentially as previously described [62] with the following modifications. Asynchronously growing cells were cultured in 15 cm dishes to ~3 × $10^7$ cells to 70-80 % confluency and labeled with EdU (10 µM, 20 min). Cells were either collected immediately (pulse) or washed with medium containing thymidine (10 µM) and either collected immediately (pulse) or incubated for 4 h (chase). Cells were fixed with 1 % paraformaldehyde in PBS for 10 min at room temperature, and fixation was quenched with 0.25 M glycine and washed with PBS. Cells were collected by scraping and centrifugation and permeabilized with 0.25 % Triton X-100 for 30 min, scraped into 15 mL tubes, and centrifuged for 5 min at 2000 rpm at 4 °C.

Pellets were washed twice with PBS and subjected to a click reaction in PBS containing 10 µM biotin–picolyl azide, 2 mM $CuSO_4$, 2 mM THPTA, and 10 mM sodium ascorbate for 25 min. Cells were centrifuged (5 min, 2000 rpm, 4 °C), washed once with PBS containing 0.5 % BSA and twice with PBS, and lysed in RIPA buffer (100 mM Tris-HCl pH 7.5, 150 mM NaCl, 1 % NP-40, 0.1 % SDS, 0.5 % sodium deoxycholate) supplemented with cOmplete™ protease inhibitor (Roche). Lysates were sonicated in 15 mL polystyrene Falcon tubes supplied with 300 µL of Bioruptor beads using a Diagenode Bioruptor Pico (20 cycles, 30 s ON / 30 s OFF) and clarified by centrifugation (10 min, max speed, 4 °C). An aliquot (100 µL) of each sample was collected as input. The remaining lysate was incubated overnight at 4 °C with 45 µL MyOne streptavidin beads. Beads were pelleted (3 min, 1800 g, RT), and the supernatant was retained as the unbound fraction. Beads were washed three times with RIPA buffer, resuspended in 65 µL RIPA, and analyzed by western blot or by LC-MS/MS.

**Sample preparation for proteomics analysis**

iPOND samples were prepared using the SP3 protocol for LC-MS/MS analysis as described[63]. In brief, samples eluted in 1xLSB buffer after iPOND were denatured at 95 °C for 10 min, and then alkylated with 400 mM IAA (final concentration) in the dark at room temperature for 30 min. Protein concentration was measured using EZQ™ Protein Quantitation Kit by following the manual. Then, protein samples were mixed and cleaned with SP3 beads (1: 10, protein: beads, with a minimum of 100 ug beads) and digested with lysC/trypsin mixture (1: 50, enzyme: protein). 2 % DMSO was used to elute the peptides from SP3 beads.

**LC-MS/MS analysis**

All MS data were acquired using a Thermo Scientific Vanquish Neo UHPLC system coupled to an Orbitrap Astral mass spectrometer with a Thermo Scientific Easy-spray source. An Ionopticks Aurora® Ultimate™ 25×75 XT C18 UHPLC column was used for the quantitative experiments with a 32-min gradient of from 8 to 35 % B and a 7.5-min gradient of from 35 % to 45 %, with a flow rate of 400 nL/min. Mobile phase A was 0.1% formic acid in water, and mobile phase B was 80 % acetonitrile and 0.1 % formic acid in water. The MS data were acquired in the data-independent acquisition mode (DIA) mode with a normalized collision energy of 25 % and a default charge state of 2. MS1 spectra were acquired in the Orbitrap at a resolving power of 240,000 every 0.6 s, with an automatic gain control (AGC) target of 500 %, covering the m/z range from 380 to 980 D. MS/MS spectra were acquired in the Astral analyser with sequential isolation windows of 4 m/z from 380 to 980, with an AGC target of $5.0 \times 10^4$ and 4 ms maximum injection time.

**MS data analysis**

All MS data were analysed together using DIA-NN version 2.2.0 [64]. The in-silico spectra library was generated by DIA-NN using the *Mus musculus* database downloaded from UniProt (SwissProt May 2025, with isoforms). The FDR threshold was set to 1 % for each of the respective Peptide Spectrum Match (PSM). The data was searched with the following parameters: stable modification of carbamidomethyl (C), variable modifications, acetylation (protein N terminus), with maximum of 1 missed tryptic cleavages threshold. MBR was checked and normalisation was disabled. The main output (*. parquet) from DIA-NN was further processed using the R package iq [65] to generate results for protein quantification. The results from 3 biological replicates were used for the following data processing.

**MS data processing in Perseus/ R-Studio**

Data processing was carried out in Perseus (v. 2.1.1.0) [66]. The dataset was median-normalised, and proteins identified with fewer than two unique peptides were excluded. To correct for batch effects, the HarmonizR plugin [67] was applied within Perseus using the Limma-based approach. Principal component analysis (PCA) and coefficients of variation (CV) were assessed both before and after batch correction to evaluate their impact. The dataset was subsequently filtered to retain only proteins detected across all three replicates at each time point, resulting in a final set of 7,139 proteins. Differential protein expression analysis was then performed in R using the Limma package[68]. P-values were adjusted using the Benjamini–Hochberg procedure [69] to control the false discovery rate, with an adjusted p-value threshold of $< 0.05$ considered statistically significant. Finally, volcano plots were generated in GraphPad Prism to visualise the differential expression results. Functional groups of proteins represented in the heatmap were selected based on the Gene Ontology database[70,71] and the STRING database[72]. Heatmaps were generated using R.

**Immunofluorescence**

Immunostaining was performed as described previously[10]. To analyze chromatin-bound proteins, cells were pre-extracted on ice with ice-cold PBS supplemented with 0.2 % Triton X-100 for 2 min. Cells were then fixed with 4 % buffered formaldehyde for 12 min at room temperature. Antibodies were diluted in DMEM (high glucose, GlutaMAX) supplemented with 10 % FBS, and incubations were performed at room temperature in a humidified chamber. Primary antibodies were incubated for 1 h, and secondary antibodies for 30 min at room temperature. Cells were washed with PBS and PBS supplemented with 0.2 % Tween-20 (Sigma-Aldrich). Coverslips were rinsed twice in distilled water, briefly dried on 3-mm filter paper, and mounted in 4.5 µl Mowiol-based mounting medium containing Mowiol 4-88 (Calbiochem), glycerol, and Tris-HCl (pH 8.5).

**Quantitative-image-based cytometry (QIBC)**

QIBC was performed as described previously[10]. In brief, high-resolution images were acquired using a ScanR inverted high-content microscope (Olympus) equipped with wide-field optics, a 20× XAPO air objective (NA 0.8), fast excitation and emission filter wheels for DAPI, FITC, Cy3, and Cy5 wavelengths, a Spectra X LED illumination system (Lumencor), and a digital monochrome Hamamatsu ORCA-Fusion camera operating in 16-bit mode. This setup yielded a spatial resolution of approximately 320 nm per pixel at 20× magnification with binning set to 1. Approximately 100 images were acquired per condition using automated ScanR acquisition software (Olympus, v. 3.6), corresponding to more than 5,000 cells.

Image acquisition parameters were adjusted to ensure non-saturated conditions within the 16-bit dynamic range, and identical settings were maintained for all samples within a given experiment. A dynamic background correction was applied before image segmentation and analysis using ScanR analysis software. Nuclear DAPI signals were used to generate an intensity-threshold mask to identify individual nuclei as primary objects. Following nuclear segmentation, pixel intensities in different channels were quantified for each nucleus or nuclear foci using predefined parameters (mean and total intensities, foci number, and foci intensities) as well as calculated parameters (e.g., sum of foci intensity per nucleus). The resulting values were exported and further analyzed using TIBCO Spotfire (v. 12.4). Comparable numbers of cells were analyzed across conditions (4,000–10,000 cells per sample). The TIBCO Spotfire software was used to generate color-coded scatter plots and calculate absolute, median, and average values for cell populations across experimental conditions. Low x-axis jittering (random displacement of objects along the x-axis) was applied to visualize overlapping cells.

**Proximity Ligation Assays (PLA)**

PLA was performed as described previously[10,73] with minor modifications. Briefly, cells were seeded on coverslips and subjected to UHRF1 gene depletion using siRNA (s26555). Following incubation, cells were fixed with methanol for 15 min and washed three times with chilled PBS. Cells were then blocked for 1 h in DMEM containing 10 % FBS and incubated with primary antibodies in a humidified chamber for 1 h at room temperature. Secondary antibody PLA probes were incubated for 1 h at 37 °C in a humidified chamber.

In situ proximity ligation was performed using a Duolink Detection Kit (Sigma-Aldrich), including ligation for 30 min followed by rolling-circle amplification for 100 min according to the manufacturer's instructions. Nuclei were stained with DAPI. Cells were washed with PBS and PBS containing 0.1 % Tween-20 after each incubation step. Nuclear PLA foci were imaged using a ScanR inverted microscope and processed by QIBC analysis. At least 8,000 cells per condition were analyzed in each experiment.

### Extended data figures:

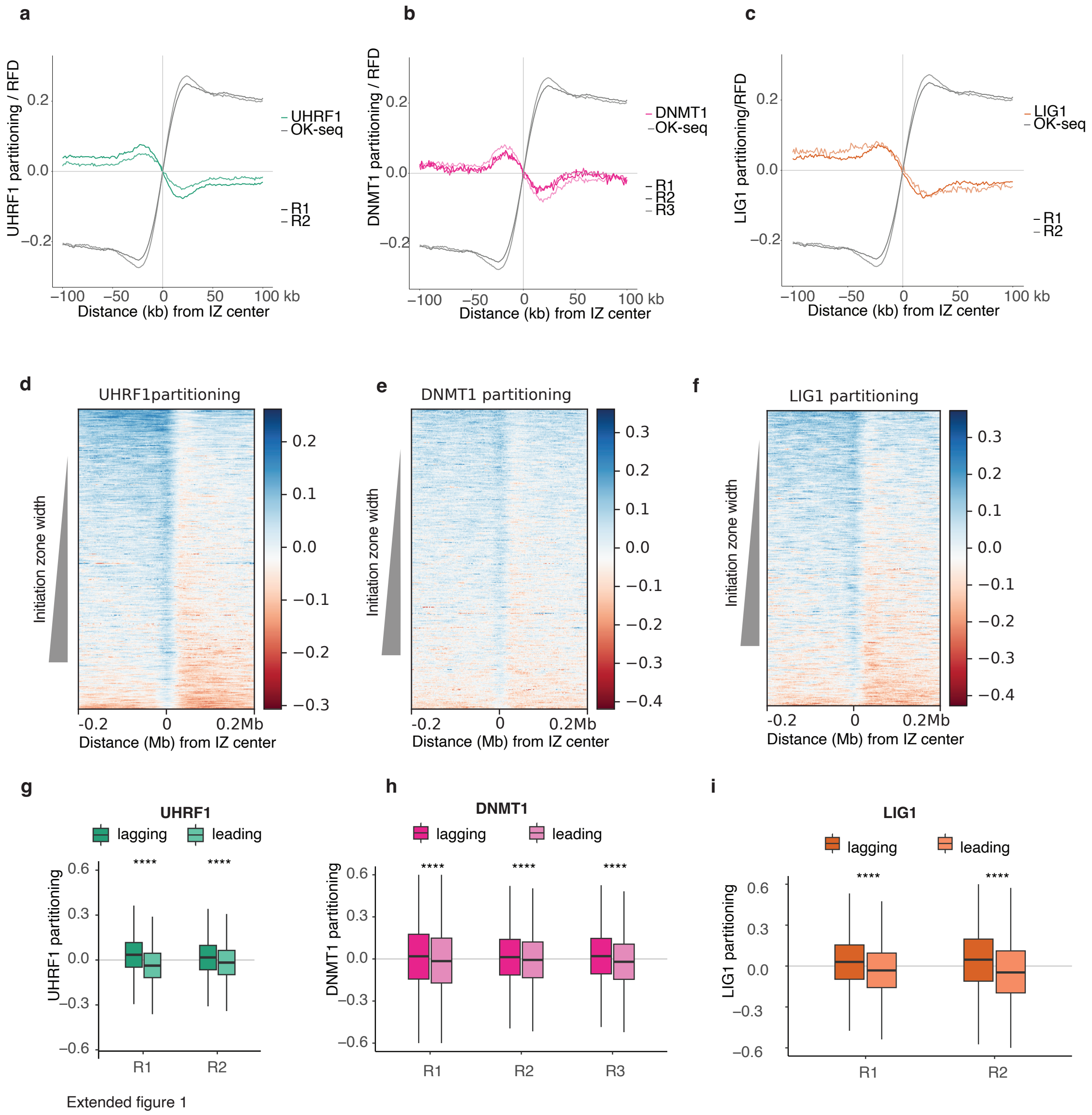


**Extended data. Fig. 1. DNMT1, UHRF1, and LIG1 show genome-wide preference towards the lagging strands.**

**(a, b, c)** Average profiles of RFD and partitioning of UHRF1 (**a**), DNMT1 (**b**), and LIG1 (**c**) around initiation zones, n=4559. Independent biological replicates are shown in different color shades.

**(d, e, f)** Heatmaps of UHRF1 **(d)**, DNMT1 **(e)**, and LIG1 **(f)** partitioning centered at initiation zones, and ordered by IZ size in descending order (Replicates 1, 3, and 1 are shown for UHRF1, DNMT1, and LIG1, respectively.)

**(g, h, i)** Boxplots of UHRF1 **(g)**, DNMT1 **(h)**, and LIG1 **(i)** partitioning values at downstream (leading, light shaded) and upstream (lagging, dark shaded) edges of initiation zones with significant partition difference in each replicate (paired Wilcoxon signed-rank test, n.s, not significant; *, p <0.05; **, p<0.01; ***, p<0.001; ****, p<0. 0001)

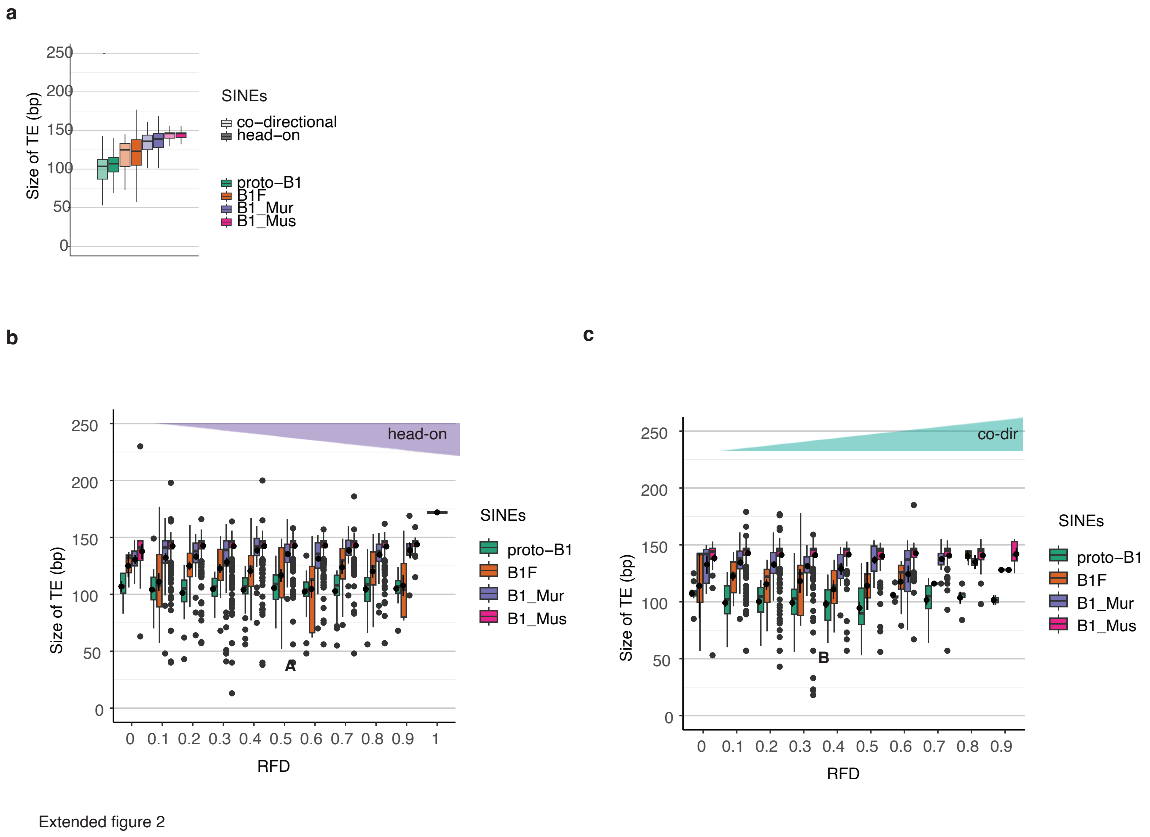


**Extended data. Fig. 2. Analysis of SINE B1 element size relative to replication fork progression in the mouse genome.**

**(a)** Boxplots of the length (bp) of distinct SINE B1 oriented co-directionally (light shaded) or head-on (dark shaded) to DNA replication forks.

**(b, c)** Boxplots of the length (bp) of distinct SINE families oriented head-on **(b)** and co-directionally **(c)** to DNA replication forks, categorized by mean RFD modal values. Higher RFD mode values indicate a greater degree of head-on (or co-directional) orientation.

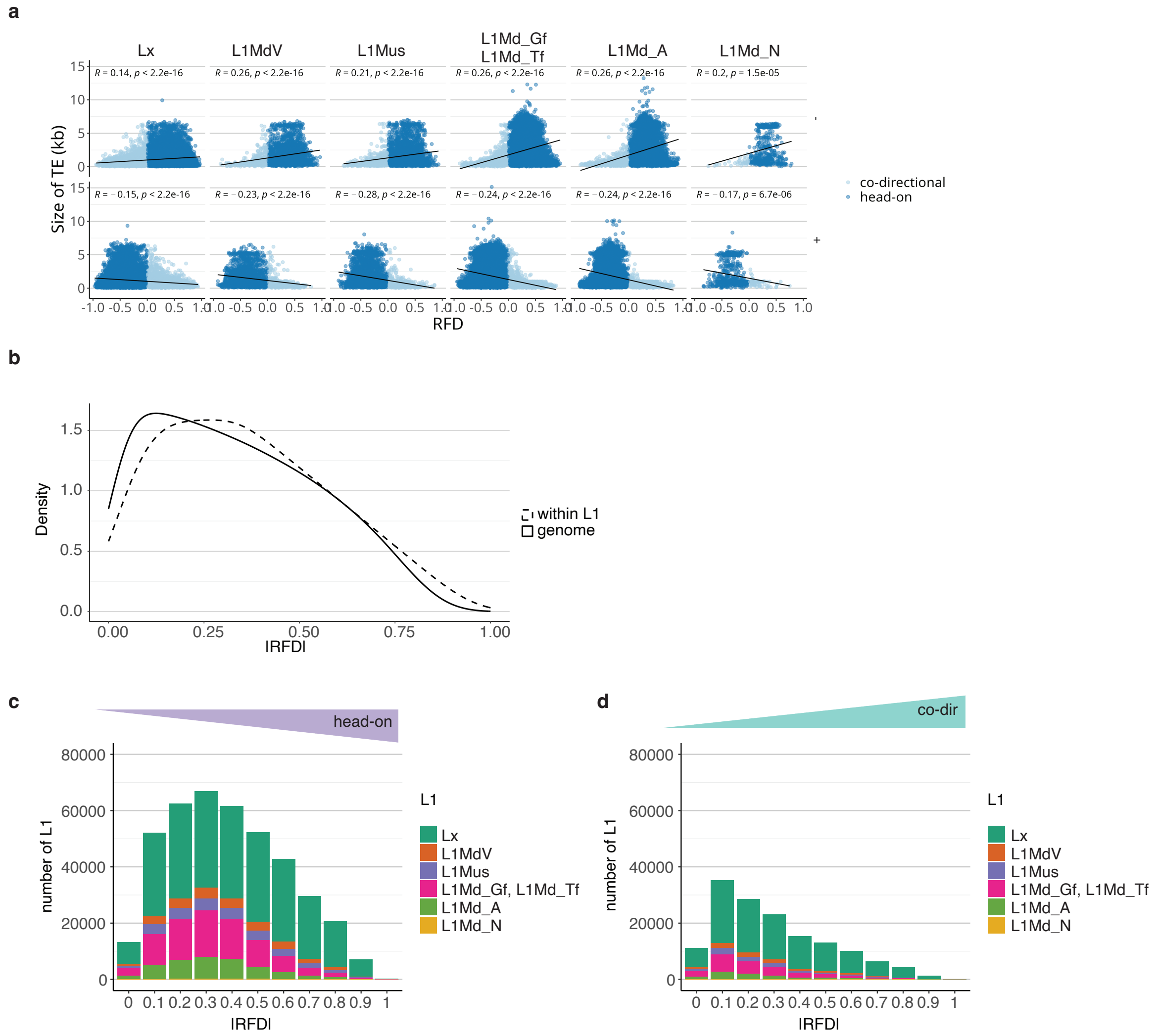


Extended figure 3

**Extended data. Fig. 3. Lagging-strand vulnerability to L1s increases over evolutionary time.**

(**a**) Scatterplots of the sizes (kb) of distinct categories of L1 versus RFD values for head-on (dark-blue) and co-directional (light-blue) for plus-strand (top) and minus-strand-encoded (bottom) L1s. Spearman correlation index and p-value are indicated on top.

(**b**) Density plots of the genomic distribution of RFD values genome-wide (plain line) and within regions containing L1s (dashed line).

(**c-d**) Bar plots of the absolute number of distinct L1 families oriented head-on **(c)** and co-directionally **(d)** to DNA replication forks, categorized by RFD modal values.

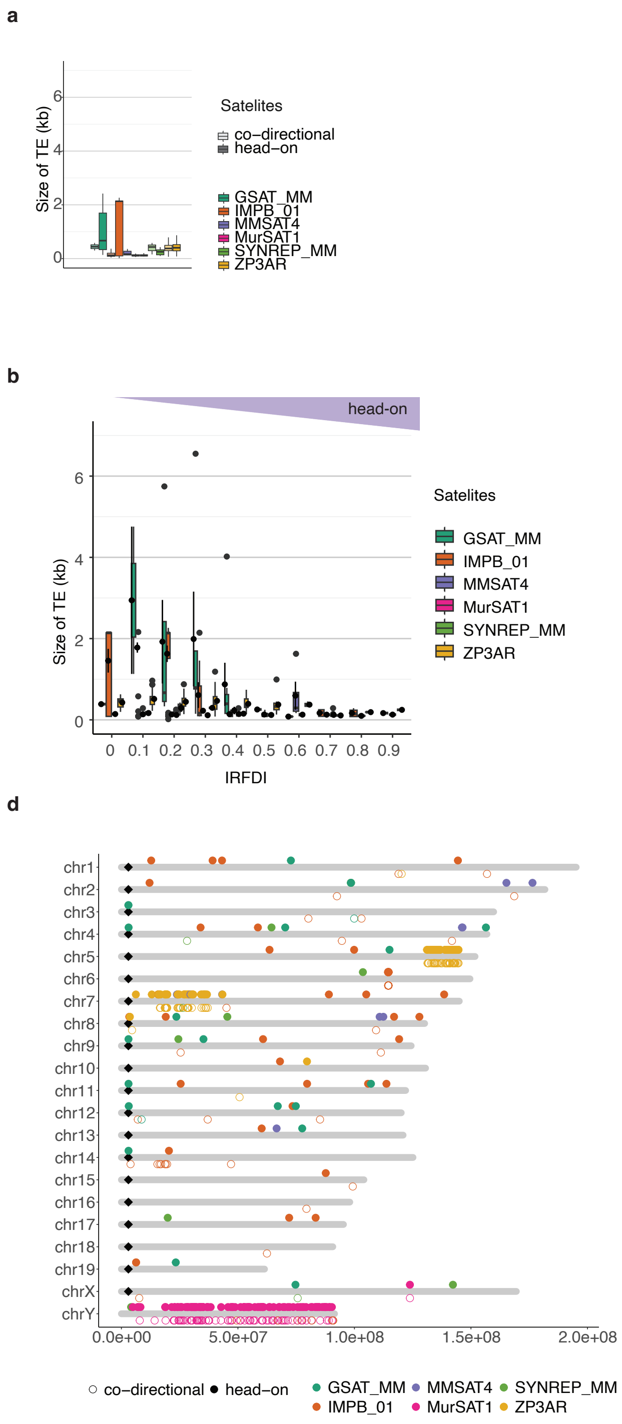


Extended figure 4

**Extended data. Fig. 4. Lagging-strand replication promotes expansion of satellites in the mouse genome.**

**(a)** Boxplots of the lengths (kb) of distinct mapped satellite repeats oriented co-directionally (light shaded) or head-on (dark shaded) to DNA replication forks.

**(b, c)** Boxplots showing the lengths (kb) of distinct mapped satellite repeat families oriented head-on **(b)** or co-directionally **(c)** to DNA replication forks, categorized by mean RFD modal values. Higher RFD mode values indicate a greater degree of head-on (or co-directional) orientation.
**(d)** Chromosomal location of mapped head-on and co-directional satellite repeats. Color shade shows distinct classes, filled circles represent head-on, and empty circles represent co-directional repeats. Black diamonds indicate centromeric positions.

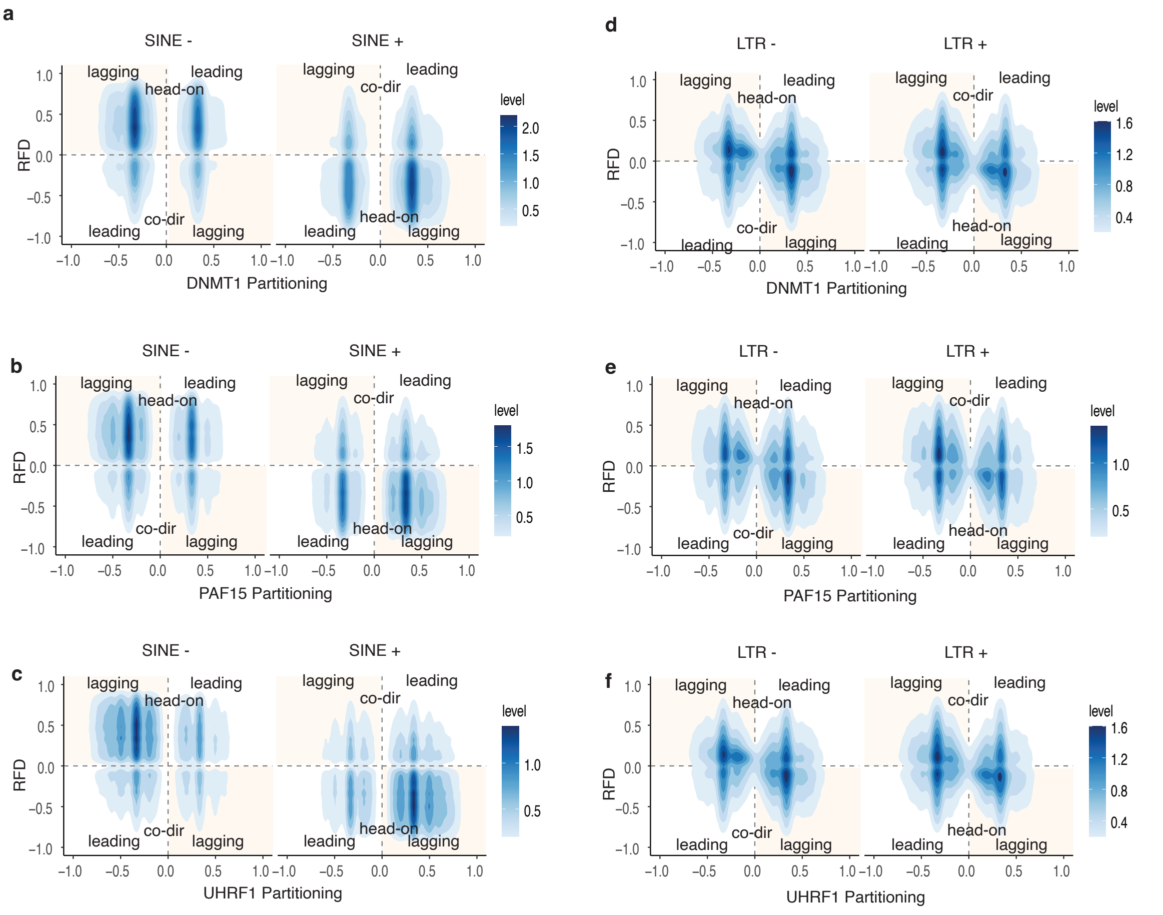


Extended figure 5

**Extended data. Fig. 5. Strand-specific enrichment of DNA methylation maintenance proteins at SINE and LTR regions during DNA replication.**

**(a, b, c)** Density plot of partitioning of DNMT1 **(a)**, PAF15 **(b)**, and UHRF1 **(c)** within leading and lagging strand replicated copies of head-on and co-oriented SINEs.

**(d, e, f)** Density plot of partitioning of DNMT1 **(d)**, PAF15 **(e)**, and UHRF1 **(f)** within leading and lagging strand replicated copies of head-on and co-directionally oriented LTRs.

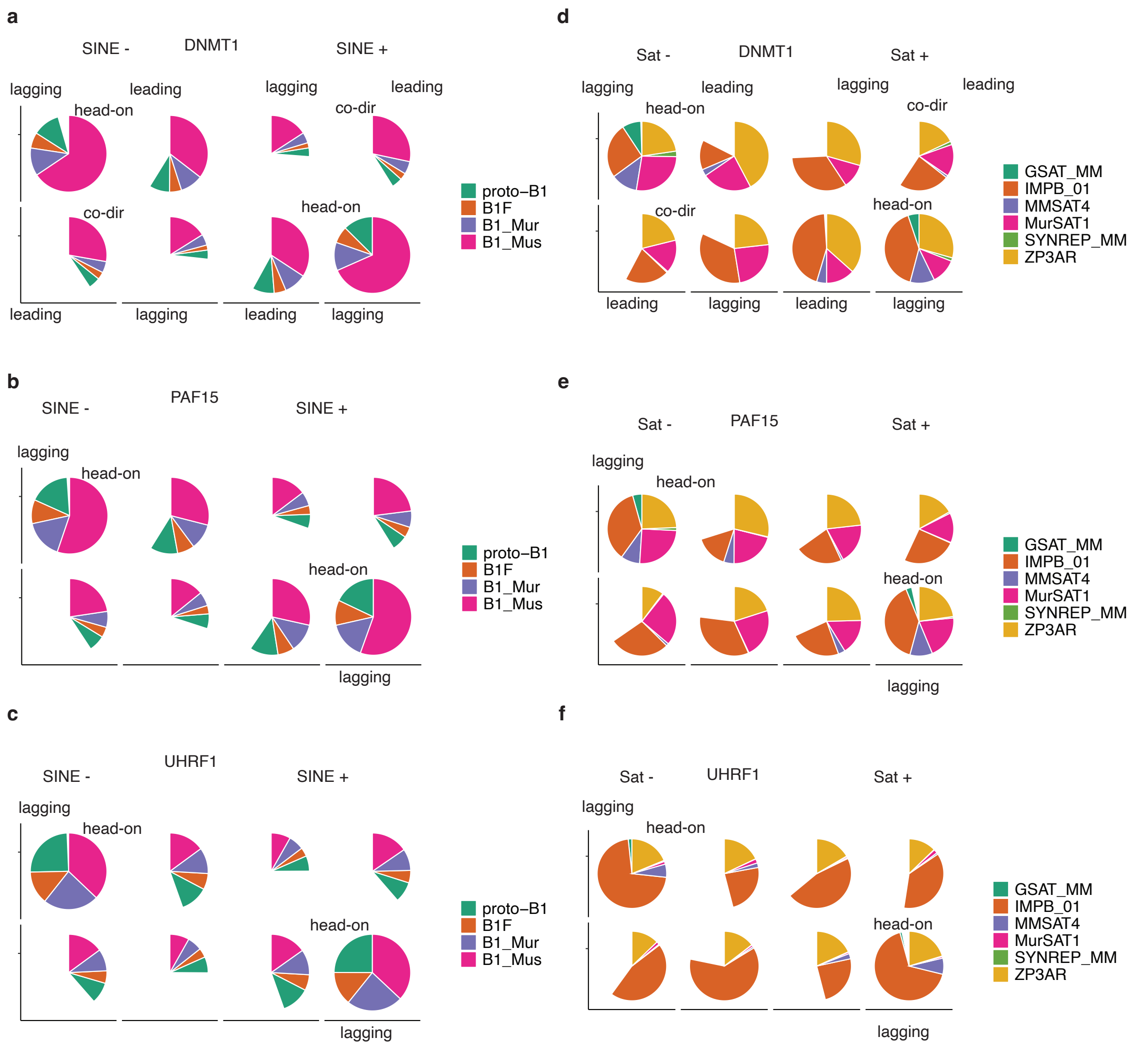


Extended figure 6

**Extended data. Fig. 6. Genomic coverage of distinct classes of SINEs and satellite repeats relative to replication mechanism and fork orientation.**

**(a, b, c)** Pie charts representing the coverage of distinct classes of newly-replicated copies of SINEs, separated into different categories of leading and lagging replication mechanisms and relative fork orientation with DNMT1 **(a)**, PAF15 **(b)**, and UHRF1 **(c)**

**(d, e, f)** Pie charts representing the coverage of distinct classes of satellites, separated into different categories of leading and lagging replication mechanisms and relative fork orientation with DNMT1 **(d)**, PAF15 **(e)**, and UHRF1 **(f)**.

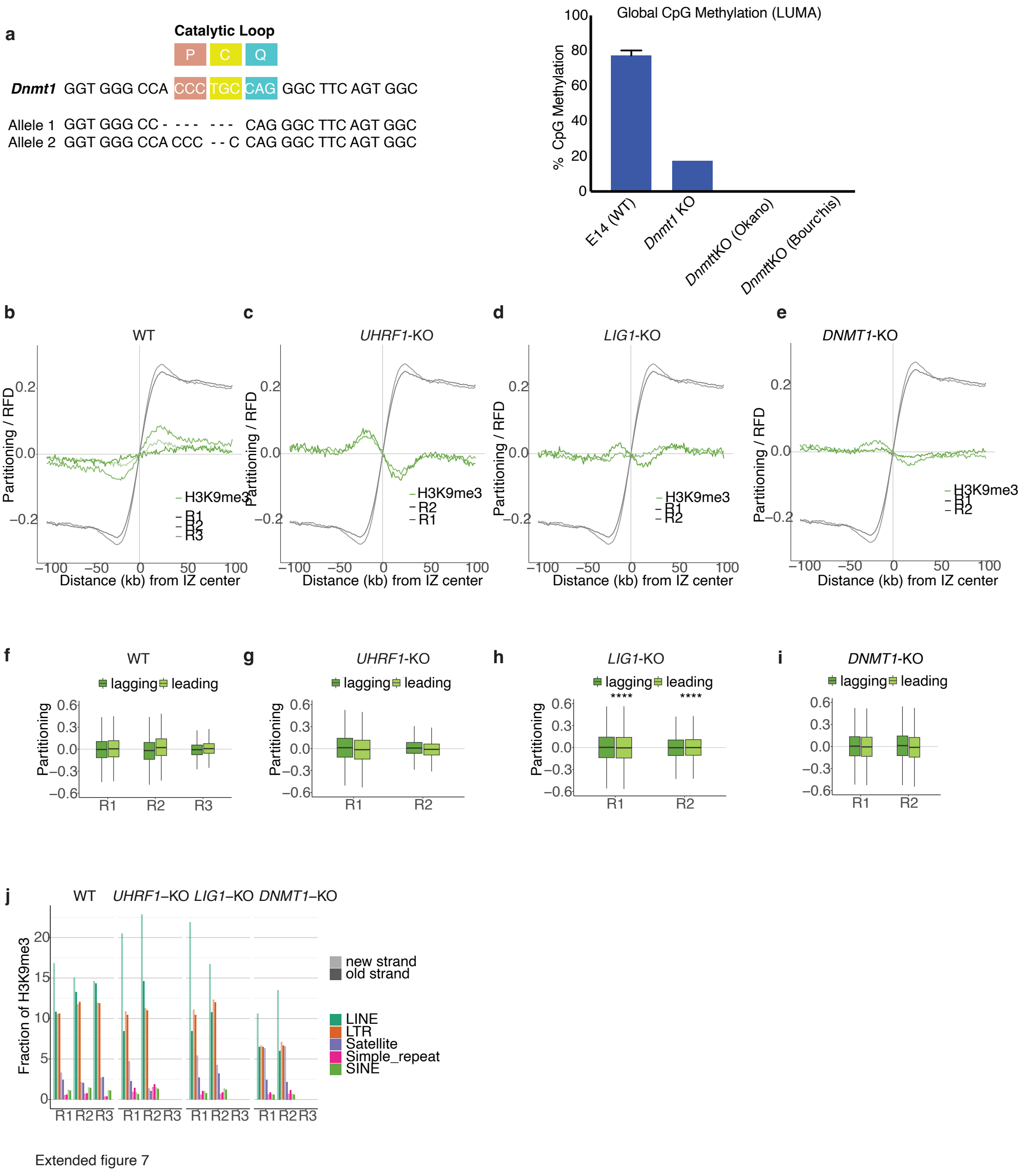


**Extended data. Fig. 7. H3K9me3 compensates for the absence of the replication-coupled DNA methylation maintenance.**

**(a)** Construction and validation of DNMT1-catalytic mutant. Left: Sanger sequencing of mutated alleles, Right: LUMA measurement of DNA methylation levels.

**(b-e)** Average profiles of RFD and H3K9me3 partitioning around initiation zones, n=4559, in wild type (**b**), *UHRF1-KO* (**c**)*, DNMT1-KO* (**d**)*,* and *LIG1-KO* (**e**) mutants. Independent biological replicates are shown in different color shades.

**(f- i)** Boxplot of H3K9me3 partitioning values at downstream (leading, light shaded) and upstream (lagging, dark shaded) edges of initiation zones with significant partition difference in each replicate of WT (**f**), *UHRF1-KO* (**g**), *DNMT1-KO* (**h**), and *LIG1-KO* (**i**) mutants (paired Wilcoxon signed-rank test, n.s, not significant; *, p <0.05; **, p <0.01; ***, p<0.001; ****, p<0.0001)

**(j)** Fraction of reads of H3K9me3 in wild type and the mutants mapped within distinct classes of TE after DNA replication, sister chromatids are annotated with respect to replicated TE orientation, with each containing either the parental (pre-existing) or newly synthesized DNA strand as the coding strand.

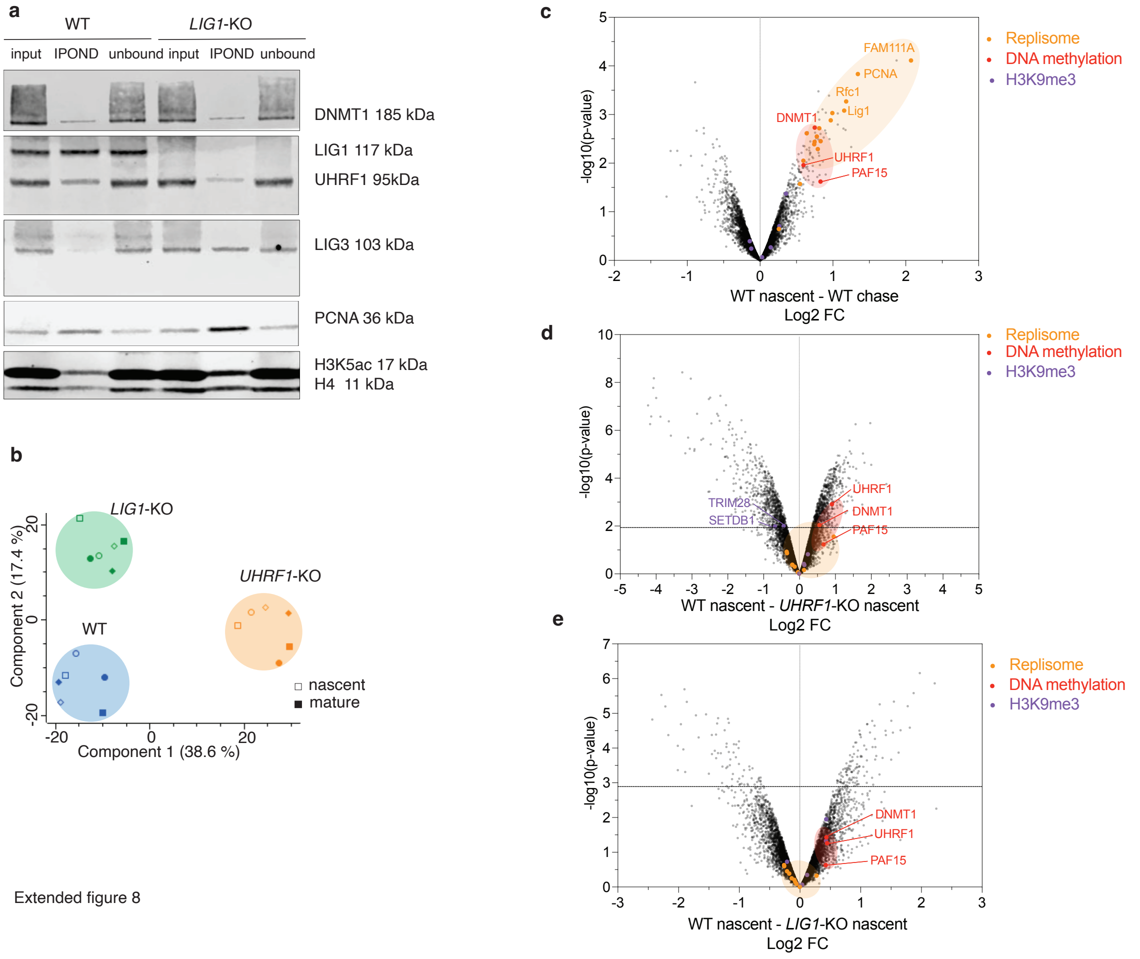


Extended figure 8

**Extended data. Fig. 8. LIG1 and UHRF1 modulate silencing mechanisms at the fork.**

**(a)** iPOND western blot comparing wild type and *LIG1-KO* mutants.

**(b)** Principal component analysis of IPOND mass-spectrometry comparing wild type and *UHRF1*-KO and *LIG1*-KO conditions.

**(c-e)** Volcano plots showing the log2 fold changes comparing protein enrichments of wild-type nascent (20 min pulse) versus mature (4 hours chase) **(c);** wild-type nascent versus *UHRF1*-KO nascent **(d)**; and wild-type nascent versus *LIG1*-KO nascent **(e).** Identified proteins related to DNA replication (orange), DNA methylation (red), and H3K9me3 (purple). The dotted line shows the threshold for statistical significance (adjusted p-value = 0.05).

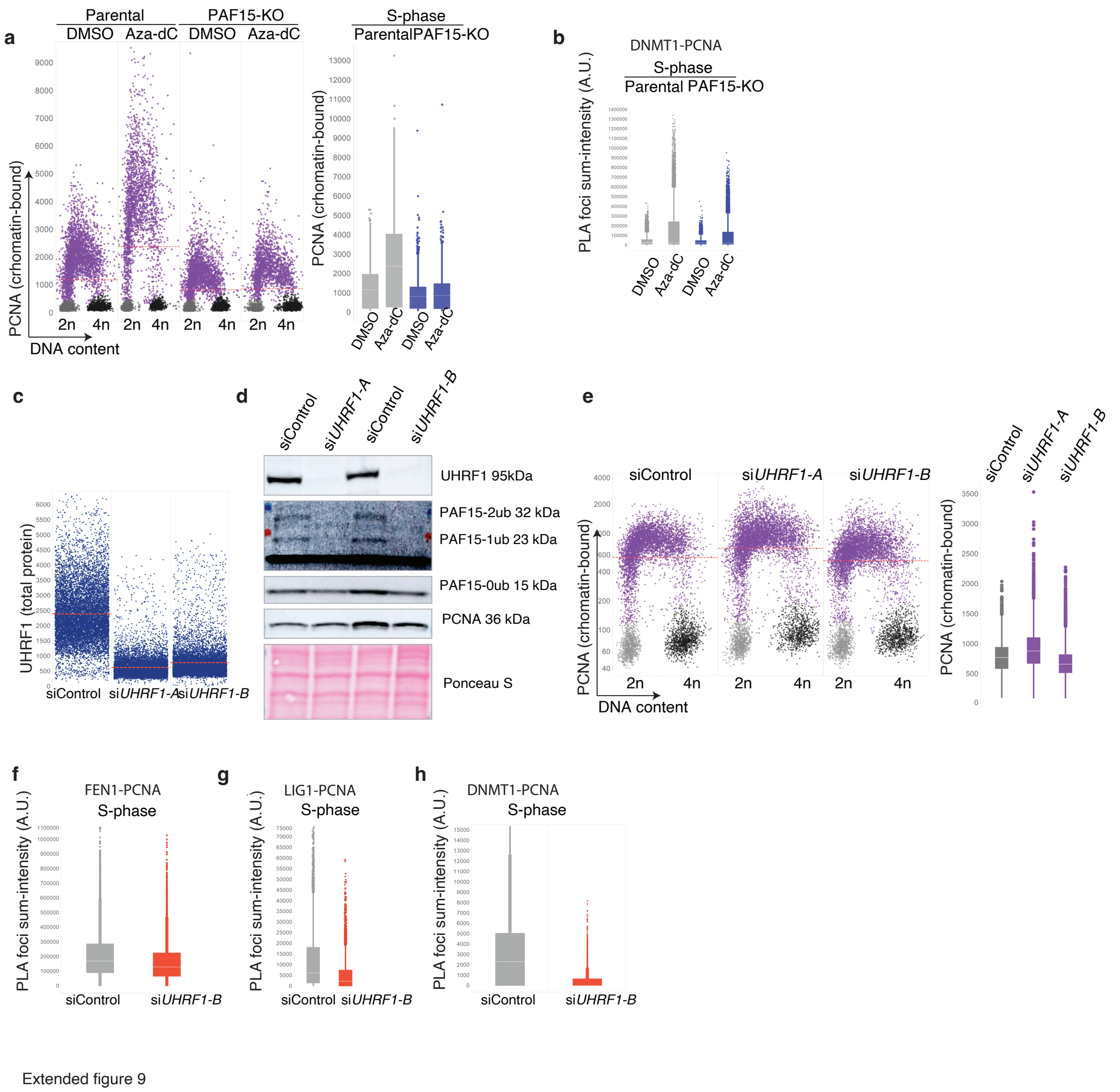


**Extended data. Fig. 9. DNMT1-UHRF1 regulates PCNA-mediated turnover at the lagging strand.**

**(a)** Right: QIBC analysis of chromatin-bound PCNA is shown as a scatter plot following decitabine (Aza-dC) treatment in U2OS parental and *PAF15*-KO (~5,000 cells analyzed per condition). DNA content: 2N (G1 phase), 4N (G2). Left: Box plot quantification of the summed intensity of PCNA in S-phase cells.

**(b)** Box plot quantification of the summed intensity of DNMT1-PCNA PLA foci in parental and *PAF15*-KO U2OS cells upon Aza-5C treatment (related to Fig. 5A).

**(c)** QIBC analysis of UHRF1 total protein intensity in U2OS parental and UHRF1-depleted cells.

**(d)** Western blot of whole cell extracts in U2OS parental and UHRF1-depleted cells.

**(e)** Right: QIBC analysis of chromatin-bound PCNA in U2OS parental and UHRF1-depleted cells shown as a scatter plot. (~5,000 cells analyzed per condition). DNA content: 2N (G1 phase), 4N (G2). Left: Box plot quantification of the summed intensity of PCNA in S-phase cells.

**(f-h)** Box plot quantification of the summed intensity of FEN1-PCNA (**f**); LIG1-PCNA (**g**), and DNMT1-PCNA **(h)** PLA foci in S-phase U2OS parental and after UHRF1 siRNA-mediated depletion cells (related to **Fig. 5c**).

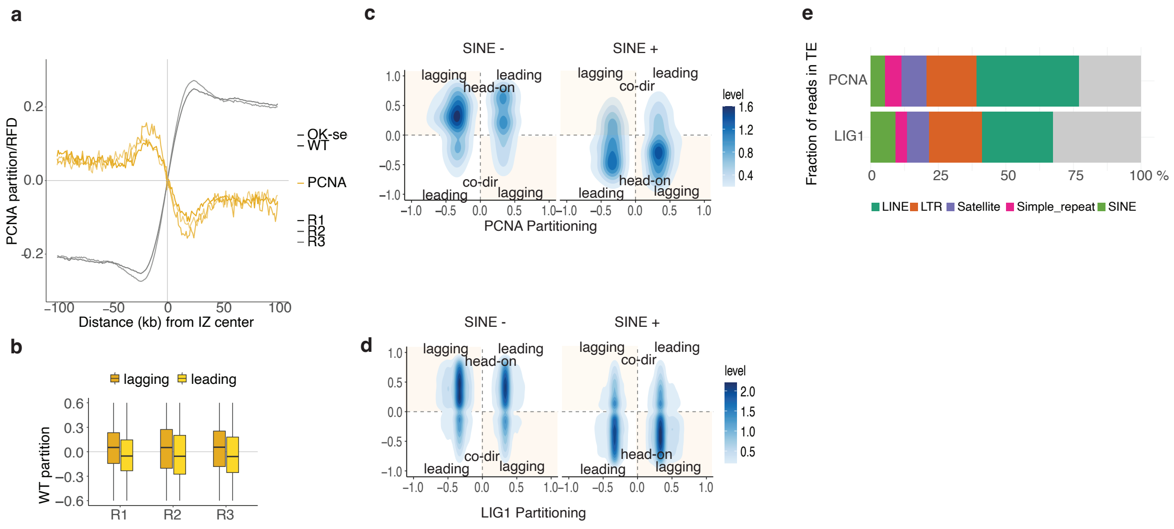


Extended figure 10

**Extended data. Fig. 10. Replication-coupled TE silencing leads to retention of replication factors PCNA and LIG1 at the TEs.**

**(a)** Average profile of RFD and PCNA partitioning around initiation zones, n= 4559. Independent biological replicates are shown in different color shades.

**(b)** Boxplot of PCNA partitioning values at downstream (leading, light shaded) and upstream (lagging, dark shaded) edges of initiation zones with significant partition difference in each replicate.

**(c, d)** Density plot of partitioning of PCNA (**c**) and LIG1 (**d**) within leading and lagging strand replicated copies of head-on and co-oriented SINEs.

**(e)** Fraction of reads of PCNA and LIG1 in transposable elements (average of biological replicates).

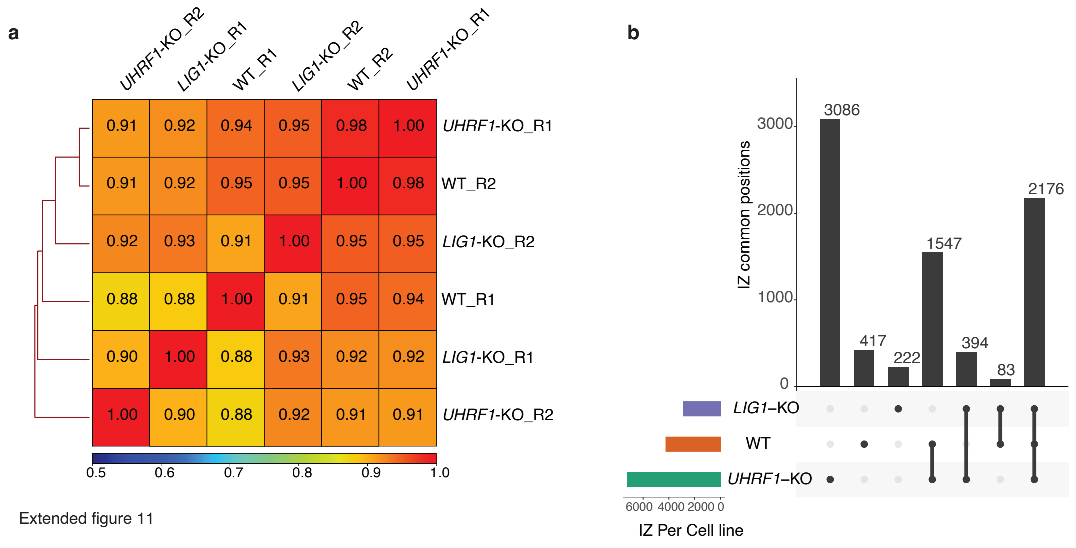


**Extended data. Fig. 11. *UHRF1*-KO and *LIG1*-KO mutations do not globally alter the DNA replication program.**

**(a)** Spearman correlation heatmap of RFD profiles.

**(b)** Upset plot of overlap of replication initiation zones between wild-type and *UHRF1*-KO and *LIG1*-KO mutants.

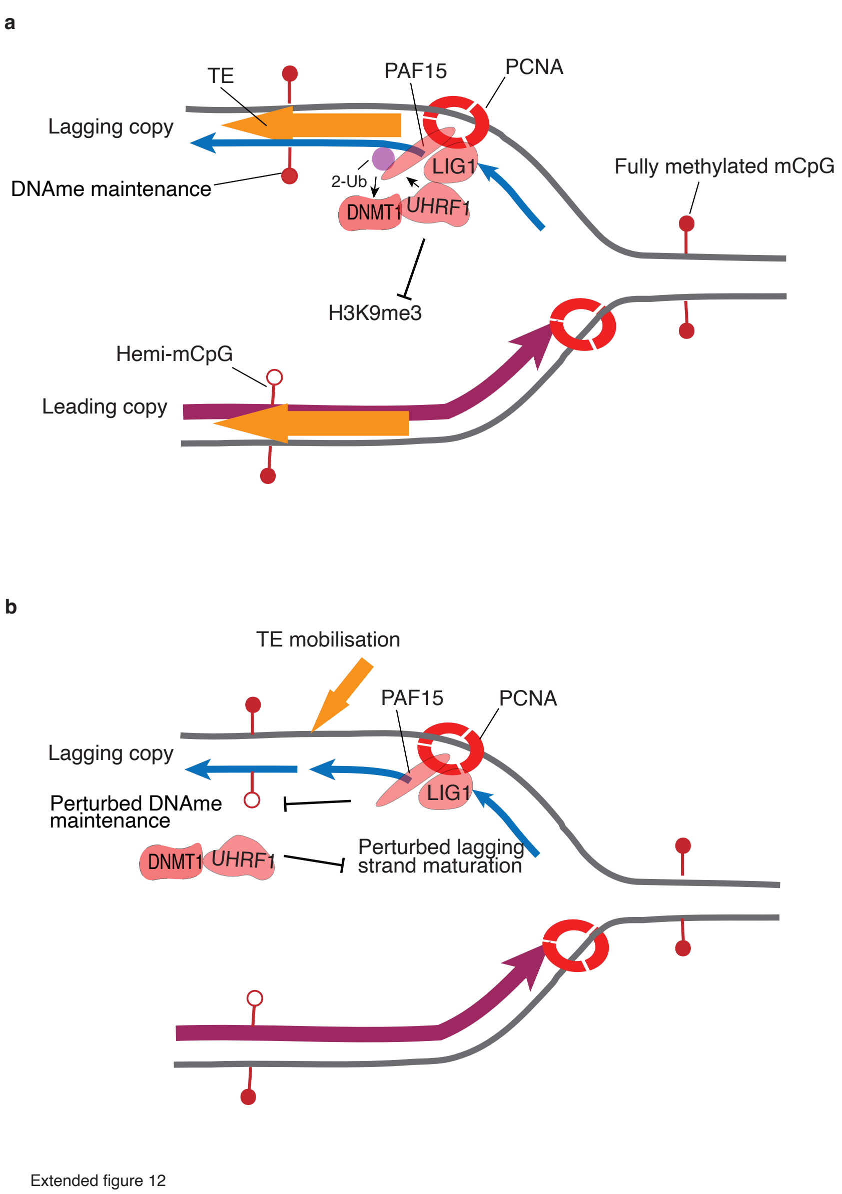


**Extended data. Fig. 12. Model: a.** Scheme representing functional coupling of the lagging strand maturation and DNA methylation maintenance through PCNA-PAF15 at regions containing TEs. The lagging strand PCNA-PAF15 interaction promotes recruitment of DNA methylation maintenance factors DNMT1 and UHRF1 at TEs. **b**. Altered DNA methylation maintenance impairs lagging-strand maturation and, thereby, may promote TE insertions at the lagging strand.
